# Contrasting evolutionary forces of specialization and admixture underlie the genomic and phenotypic diversity of *Yarrowia lipolytica*

**DOI:** 10.1101/2025.11.28.690175

**Authors:** Sergio Izquierdo-Gea, Javier Vicente, Tomás A. Peña, Pablo Villarreal, Francisco A. Cubillos, Cécile Neuvéglise, Jorge Barriuso, Ignacio Belda, Javier Ruiz

**Author notes:** Corresponding authors: Javier Ruiz; Ignacio Belda; Jorge Barriuso.

## Abstract

Microbial diversity emerges from evolutionary processes that shape genomic and phenotypic traits in response to complex environmental pressures. Deciphering these dynamics is key to understanding microbial ecology and advancing biotechnological applications. Here, we use the yeast *Yarrowia lipolytica* to study genomic signatures of adaptation across a broad range of environments and to illustrate how population-level data can inform targeted bioprospecting for industrial traits. Whole-genome and phenotypic analyses of 126 isolates from natural and anthropogenic environments reveal a complex population structure in this species, shaped by both niche specialization and admixture events. Structured lineages exhibit ecological filtering, reduced genetic diversity, and distinct gene content, consistent with adaptation to substrates like dairy, hydrocarbons, or industrial substrates. In contrast, admixed populations display greater genetic diversity and broader phenotypic capacity, including enhanced stress tolerance and metabolic flexibility. Genome plasticity, reflected in pangenome and CNV variation, aligns with ecological origin, while trait assays link phenotypic divergence to underlying genetic variation. For instance, better growth performance on acetate, an ecological and industrially relevant trait, is associated with hydrocarbon-adapted strains and likely linked to variation in regulatory regions of acetate metabolism genes. Together, our results reflect how divergent evolutionary trajectories—ranging from ecological specialization to genomic plasticity through admixture— underpin the species’ ecological success and provide a framework for harnessing its natural diversity in microbial bioprospecting.

## INTRODUCTION

Microbes thrive across an extraordinary range of environments, from nutrient-rich substrates to extreme, resource-limited ecosystems. Understanding how microbial diversity emerges and persists across natural and anthropogenic environments remains a central challenge in evolutionary biology, with significant implications for ecology, biotechnology, and microbial bioprospecting. In particular, the extensive genomic and functional variation that underlies microbial diversity offers unique insight into the evolutionary mechanisms that enable adaptation to diverse niches (Cohan & Koeppel, 2008; Peter & Schacherer, 2016). Among eukaryotic microbes, yeasts represent an especially tractable system for studying these dynamics due to their short generation times, ease of cultivation, and well-characterized genomes. Specifically, yeasts of the subphylum *Saccharomycotina* have repeatedly adapted to both natural and anthropogenic environments, providing a valuable framework to explore how environmental pressures drive diversification, specialization and genome evolution (Gonçalves et al., 2016; Harrison et al., 2024; Opulente et al., 2024; Starmer & Lachance, 2011).

While the model yeast *Saccharomyces cerevisiae* has received considerable attention—especially for its domestication history and adaptive divergence in anthropogenic niches (Almeida et al., 2015; Bigey et al., 2021; Steensels et al., 2019)—many other species exhibit similarly broad ecological distributions. Comparative studies in wine, beer, and other fermentation ecosystems have revealed clear genomic signatures of niche-specific adaptation, highlighting the power of population-level analyses to illuminate the genomic basis of microbial specialization (Almeida et al., 2014; Albertin et al., 2016; Eberlein et al., 2021; Vicente et al., 2024). Yet beyond the realm of fermented beverages, the adaptive landscapes of yeasts remain underexplored, with few exceptions regarding dairy products and clinical settings (Bennetot et al., 2023; Friedrich et al., 2023; Sephton-Clark et al., 2022; Wang et al., 2024). This narrow focus likely imposes a strong bias on our understanding of yeast evolutionary dynamics and constrains our ability to generalize across broader phylogenetic scales.

*Yarrowia lipolytica* stands out as an ideal model for addressing this gap. This species combines features rarely found together: a cosmopolitan distribution across diverse natural and anthropogenic habitats, a high metabolic versatility, and a recent but well-established history of industrial application (Madzak, 2021). It is frequently isolated from marine and terrestrial environments, insect hosts, animal-derived substrates, fermented foods, and hydrocarbon-rich industrial settings, highlighting its ecological plasticity and ability to colonize challenging habitats (Madzak, 2021; Mamaev & Zvyagilskaya, 2021). Despite this unique confluence of ecological breadth and biotechnological relevance, evolutionary and ecological studies on *Y. lipolytica* remain sparse. Research has disproportionately focused on a narrow set of laboratory strains—most notably W29 and its derivatives—selected for traits such as lipid accumulation, organic acid production, and stress tolerance (Liu et al., 2015; Madzak, 2018). While a few wild isolates (e.g., A-101, H222) have been examined for phenotypic diversity, broader surveys capturing the species’ full ecological and genetic range are still scarce (Quarterman et al., 2017; Bigey et al., 2023). Intriguingly, early comparative genomics in this species have revealed substantial phenotypic divergence and low genetic diversity in the absence of clear population structure linked to geographic or ecological origin, raising fundamental questions about the evolutionary forces driving this disparity (Bigey et al., 2023). As such, *Y. lipolytica* offers a powerful, underutilized system for investigating how environmental heterogeneity, anthropogenic selection, and genome evolution intersect to shape microbial diversity.

In this work, we leverage a broad collection of 126 *Y. lipolytica* isolates to understand the evolutionary and ecological processes that underpin its present-day diversity. Our analyses uncover a complex population structure that is strongly associated with ecological origin and is accompanied by genome-wide variation in gene content and copy number. We identify both genetically admixed populations exhibiting broader phenotypic performance across diverse growth conditions and highly structured, ecologically specialized lineages, suggesting the coexistence of divergent evolutionary trajectories that support adaptation to human-associated environments. By linking population structure to growth performance and metabolic traits under industrially relevant conditions, we lay the groundwork for targeted strain selection and rational bioprospecting, revealing how ecological pressures in anthropogenic environments have shaped functional diversity in this emerging microbial cell factory.

## RESULTS

### Adaptation to different anthropogenic environments drives the population structure of *Y. lipolytica*

To explore the genomic diversity and adaptive potential of *Y. lipolytica*, we assembled a dataset of whole-genome sequences from 126 isolates spanning a broad range of isolation sources. The majority of isolates (70.3%) originated from anthropogenic environments, including foods, urban and industrial settings, which reflects the global strain distribution in culture collections (Table S1).

Population structure analyses of the species, based on 207,153 genome-wide SNPs, revealed six genetic groups (Fig. 1A–B). These clusters were consistently supported by complementary approaches, including principal component analysis, haplotype-based clustering and SNPs density maps (Fig. S1, S2A-B, S3), and showed significant association with both isolation source and geography (Chi-squared test, p = 4.26 × 10⁻⁸ and 7.75 × 10⁻⁹, respectively; Fig. S4A–B). Based on their predominant ecological origin and main geographical component, we designated the groups as follows: strains from plant-based industrial settings in North America (IndNA), soil and hydrocarbon-associated strains from Eurasia (Soil-Hydrocarbon, So-HC), food-and host-associated strains from Europe (Eur1/Mix), and dairy-derived strains from Europe (Eur2/Dairy). Additionally, the clonal lineage harbouring the reference strain W29 was named W29 clade (dark blue, Fig. 1A) and several admixed strains were grouped under the term Mosaic (grey, Fig. 1A). Together, phylogenetic and population structure analyses suggest an early divergence of North American strains (IndNA), followed by the diversification of the remaining Eurasian lineages adapted to distinct anthropogenic environments.

**Figure 1.**
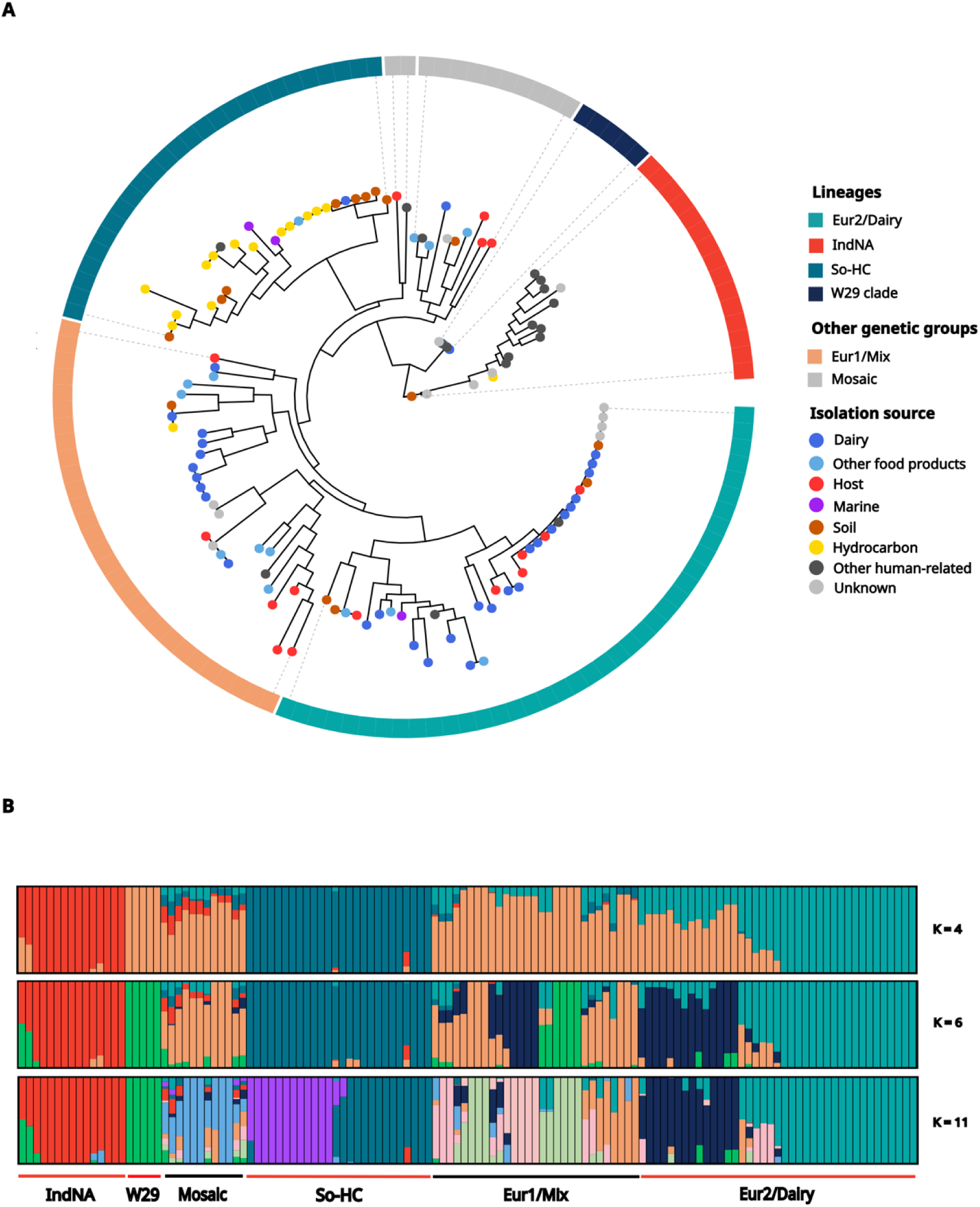
Population structure analysis of 126 *Yarrowia lipolytica* strains. (A) Maximum likelihood phylogenetic tree of the 126 *Y. lipolytica* strains. Tip colors indicate isolation source. The outer ring represents the proposed population structure model integrating ADMIXTURE, fineStructure, PCA and phylogeny. The tree is rooted with the strain closest to *Y. yakushimensis*, the nearest known species within the genus. (B) ADMIXTURE results for K = 4, 6, and 11 ancestral populations, shown in phylogenetic order. Colored bars represent ancestry components; red lines below indicate structured populations, black lines indicate admixed strains.

To better understand the genomic basis underlying the observed patterns of population structure, we performed a pangenome analysis of *Y. lipolytica*. The resulting pangenome comprises 6,581 genes, including 6,193 core genes, shared by all strains, and 139 soft-core genes (shared by >95% of strains). The accessory genome consists of 249 genes, split nearly evenly between medium-frequency (5–95% of strains, 126 genes) and rare (<5%, 123 genes) genes. The pangenome is open (Heap’s α = 0.915), meaning new genes are expected with additional genomes (Fig. S5). Notably, gene content variation in the accessory genome is shaped by population structure, which explains 21.5% of the observed variance (PERMANOVA, *p* = 0.001, Fig. S6). In parallel, we investigated copy number variants (CNVs) as an additional source of genome plasticity. We identified 4,783 CNV events affecting 2,083 core coding sequences, excluding transposons and pseudogenes (Table S2). Both CNV frequency and total genomic length differed significantly across genetic groups for gains and losses (Kruskal–Wallis test, *p* < 0.01, Fig. S7).

Importantly, the six genetic groups identified fall into two contrasting patterns. Four of these (IndNA, So-HC, W29 clade and Eur2/Dairy) showed strong concordance with phylogenetic lineages and specific isolation sources, supporting their evolutionary cohesion and likely reflecting adaptation. In contrast, the Eur1/Mix group and Mosaic strains displayed lower resolution across population structure-based analyses and more diverse isolation sources, suggesting more diffuse or reticulate evolutionary histories. These contrasting patterns highlight the coexistence of both structured, adapted lineages and more flexible admixed genetic groups, suggesting a complex evolutionary history linked to anthropogenic environments.

In particular, the admixed genetic groups (Mosaic and Eur1/Mix) illustrate a discordance between phylogeny and population structure, revealing reticulate evolutionary processes that escape bifurcating phylogenetic models. Neither group is monophyletic, yet both share consistent genomic signatures across structure-based analyses, suggesting that admixture has played a central role in shaping their genomes. These admixed populations exhibit weak population structure as evidenced by their lower pairwise genetic differentiation (Fst = 0.14-0.38; Fig. S8A) and high nucleotide diversity (π = 1.73×10⁻³ and 1.75×10⁻³; Fig. S9A), consistent with their complex genomic backgrounds. Haplotype co-ancestry analysis (Fig. S2A) reveals extensive SNP sharing between Eur1/Mix and IndNA strains, while Mosaic strains show gene flow with the W29 clade and IndNA (D-statistic: D=0.61, Z=7.65, p = 2.02×10^⁻14^; D=0.57, Z=9.22, p = 2.3×10^⁻16^, respectively) and low divergence from Eur1/Mix despite their phylogenetic separation (Fig. S8A), confirming admixture and historical connectivity across lineages (Fig. S8A).

To ensure that the observed patterns were not driven solely by the polyphyletic nature of these groups, we analyzed each monophyletic clade independently. Genetic diversity parameters remained within the same range (F_st_ = 0.08–0.30; π = 1.2–1.8×10⁻³, Fig. S8B and 9B, respectively), confirming that their high nucleotide diversity and weak structure are intrinsic features, not artifacts of polyphyly. Finally, the two groups exhibit contrasting genome dynamics: Eur1/Mix strains are enriched in gene amplifications, while Mosaic strains show the highest rates of gene loss (Fig. S7A), pointing to distinct evolutionary trajectories after admixture.

As opposed to the admixed genetic groups, more well-defined lineages (IndNA, So-HC, W29 clade, and Eur2/Dairy) exhibit lower nucleotide diversity (π = 1.7×10⁻⁵ to 9.57×10⁻⁴; Fig. S9A), consistent with genetic bottlenecks or limited standing variation. This is further supported by moderately positive Tajima’s D values in IndNA (0.75), So-HC (1.02) and the W29 clade (0.67), indicative of past demographic contractions or population structure. In contrast, Eur2/Dairy displays a near-neutral Tajima’s D (–0.02), suggesting a population close to mutation-drift equilibrium, with no strong evidence for recent demographic shifts or selective sweeps (Fig. S10).

In line with these genomic features, these lineages also harbor distinctive gene content profiles that may reflect adaptive responses to specific ecological pressures. For example, the IndNA population, one of the earliest diverging lineages, shows distinct gene amplifications, including in pathways for amino acid and cofactor biosynthesis, as well as responses to nutrient limitation. Interestingly, many of these amplified genes correspond to gene losses in other groups. The So-HC population includes a clonal European subgroup enriched for genes involved in iron acquisition, possibly reflecting adaptation to soils and hydrocarbon-contaminated environments (Fig. 1A). Eur2/Dairy stands out for its markedly low nucleotide diversity (Fig. S9A and S9C) and minimal gene content variability, consistent with clonality. It lacks lineage-specific genes, shows the shortest amplified genome length, and contains the fewest genes affected by amplifications (Fig. S7A-B). The few CNV gains it displays are mostly associated with transposon-related sequences. These findings, together with Eur2/Dairy forming the most distinct coancestry cluster (Fig. S2B), and occupying a terminal branch in the phylogeny (Fig. 1A), support its status as a recently diverged population shaped by clonal propagation and ecological constraints.

In summary, our findings uncover two contrasting evolutionary patterns across *Y. lipolytica* populations: while some groups exhibit flexible, admixed genomic backgrounds with higher genetic diversity, others show strong ecological specialization. These specialized populations are tightly associated with specific anthropic environments, suggesting adaptation to industrial or human-modified habitats. This adaptation is accompanied by directional changes in gene content and reduced genomic diversity, hallmarks of niche specialization, highlighting the species’ broader tendency toward adaptation to anthropic settings.

### Functional diversity in *Y. lipolytica* reflects genetic structure and environmental adaptation

To explore how genetic diversity translates into phenotypic variation and potential environmental adaptation, we analyzed 49 traits across strains isolated from diverse environments. These included growth and lipase activity under various environmental (temperature, pH, nitrogen availability and chemical stressors) and nutritional (different carbon and nitrogen sources) conditions, as well as protease activity and colony morphology (Table S3).

We first studied whether phenotypic variation mirrors the underlying phylogenetic relationships among strains. A significant phylogenetic signal was detected for over half of the measured traits (19 out of 35; Table S4), and pairwise phenotypic and phylogenetic distances were positively correlated (r = 0.30, p = 2.2×10⁻¹⁶, Fig. S11), indicating that evolutionary relatedness contributes to phenotypic similarity. Consistently, population structure explained 15.8% of the total phenotypic variance (PERMANOVA, *p* = 0.001), and phenotypic variability—measured via coefficients of variation— was significantly lower within populations than between them (*p* = 0.006, Fig. S12). Together, these findings support the existence of genetically encoded phenotypic differentiation across major *Y. lipolytica* clades.

Most strains failed to grow on 12 of the 28 carbon sources tested, which were excluded from further analysis (Table S3). Among the retained conditions, several exhibited binary (growth/no-growth) responses across strains, particularly growth in citrate and nitrates (as sole carbon and nitrogen sources, respectively), at high temperature (37 °C), and under certain stressors. Most other conditions supported growth and exhibited considerable quantitative variability. Notably, 22 traits significantly differed between populations (Kruskal-Wallis test, *p* < 0.05; Fig. 2A).

**Figure 2.**
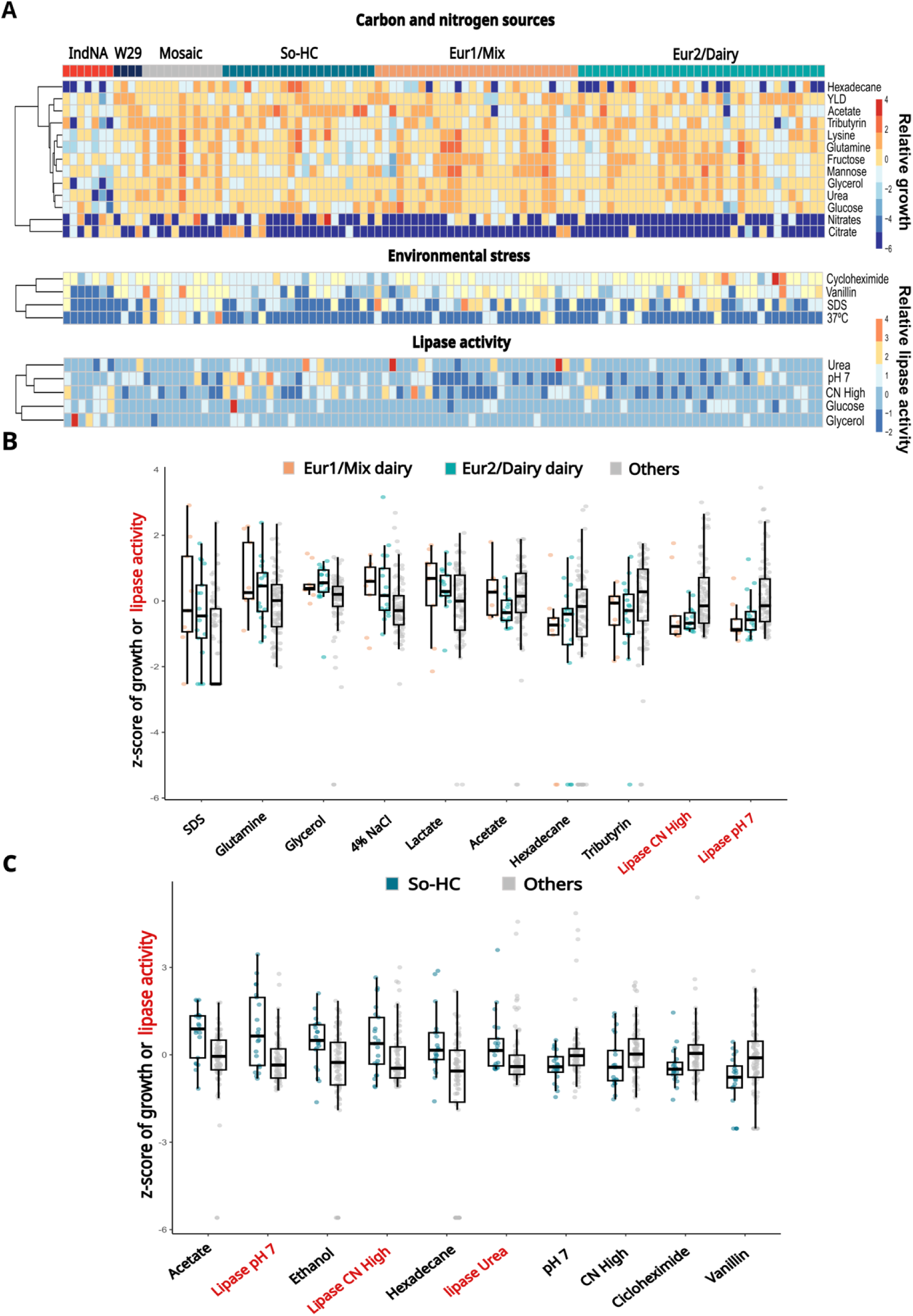
Phenotypic analysis of a subset of 105 *Y. lipolytica* strains encompassing the ecological diversity of the whole dataset. **(A)** Heatmap of conditions in which populations behave significantly differently. For carbon and nitrogen sources, the parameter represented is the z-score of OD_600_ at 48 hours of culture. In the case of environmental stress conditions, the z-score of Eff 48, previously normalised by growth in glucose, is shown. For lipase activity, the z-score of units of activity, normalized by OD600 at the measurement time, is the parameter represented. Strains are ordered according to phylogenetic relationships and population structure is annotated at the top label. **(B)** Traits indicative of adaptations in dairy environment, as evidence by statistical testing (Wilcoxon rank sum test, p < 0.05). Dot colours indicate the population, while red labels refer to lipase activity. **(C)** Traits indicative of adaptations to So-HC, as evidence by statistical testing (Wilcoxon rank sum test, p < 0.05). Dot colours indicate the population, while red labels refer to lipase activity.

Regarding strains from admixed groups, namely Mosaic and Eur1/Mix, they exhibited broad niche breadth, indicated by significantly higher growth across diverse environmental contexts (Kruskal-Wallis test, p = 9×10⁻7, Fig. S13A). Mosaic strains, in particular, showed robust growth across diverse conditions, including elevated temperatures, and consistently outperformed other groups (Fig. S13B). This may be linked to the admixture events affecting this clade may have conferred increased phenotypic robustness. Similarly, Eur1/Mix strains displayed high growth rates in most conditions, although a notable fraction was unable to utilize citrate or nitrates. (Fig. 2A). Notably, both groups predominantly formed smooth colonies, indicative of reduced invasiveness and a predominance of yeast-like morphologies (Fig. S14).

Different phenotypic responses were observed among strains from genetically structured populations. Strains from the IndNA clade exhibited overall reduced growth and marked sensitivity to SDS, vanillin, and copper. However, these traits did not clearly reflect adaptation to a particular environment, and may instead reflect a conserved or ancestral phenotypic profile (Fig. 2A). The W29 clade, largely represented by the laboratory reference strain and its derivatives, showed robust growth across conditions but lacked consistent patterns suggestive of ecological specialization.

Contrary, other genetically structured populations showed phenotypic profiles more clearly aligned with ecological specialization. Eur2/Dairy strains, associated with dairy products, exhibited consistent adaptation to milk-derived environments, including enhanced growth on lactate, NaCl, and SDS (Wilcoxon rank-sum test, *p* = 0.043, 0.008, and 0.003, respectively), and reduced lipase activity at pH 7 and under low availability of nitrogen (C/N High, Fig 2B), as well as impaired growth on tributyrin and acetate (*p* = 0.0004 for both, Fig. 2B). Some of these traits were shared with dairy strains from the Eur1/Mix clade, including more frequent absence of protease activity (Chi-squared test, *p* = 0.03, Fig. S14), reinforcing that they reflect adaptation to dairy beyond shared ancestry (Fig. 2B).

Finally, strains from soil and hydrocarbon-associated environments (So-HC) demonstrated superior performance in traits related with hydrophobic substrates. These strains showed enhanced growth on hexadecane, acetate, and ethanol, carbon sources likely prevalent in their native habitats (Fig 2C). They also exhibited increased lipase activity across multiple conditions and rough colony morphologies, suggestive of enhanced invasiveness and likely reflecting adaptive responses to fluctuating, resource-limited environments.

Together, these results show that phenotypic variation in *Y. lipolytica* recapitulates both the genomic complexity of admixed lineages and the ecological specialization of genetically cohesive populations. This phenotypic landscape highlights the role of habitat-specific selective pressures and provides a framework for informed bioprospecting in industrial and environmental contexts.

### Population-specific genomic adaptations guide trait-based bioprospecting

After identifying clear ecological specialization in Eur2/Dairy and So-HC strains, we explored the underlying genetic bases of these adaptations. To this end, we performed functional enrichment analyses focusing on population-specific gene amplifications and genes exhibiting strong allele frequency divergence (genes with higher mean F_st_).

In the Eur2/Dairy population, gene copy number gains were enriched in divalent cation transporters (e.g., zinc, copper, magnesium; Fig. S15), whereas the most differentiated genes were related to amino acid metabolism and transport (Fig. 3A), consistent with their superior growth on amino acids as nitrogen sources (Wilcoxon rank-sum test, p < 0.05). For the So-HC group, gene amplifications and differentiated genes were enriched in oxidative stress response, actin nucleation and xenobiotic transport, potentially supporting their enhanced growth on ethanol, acetate, and other hydrophobic substrates. (Fig. S16A and 3B).

**Figure 3.**
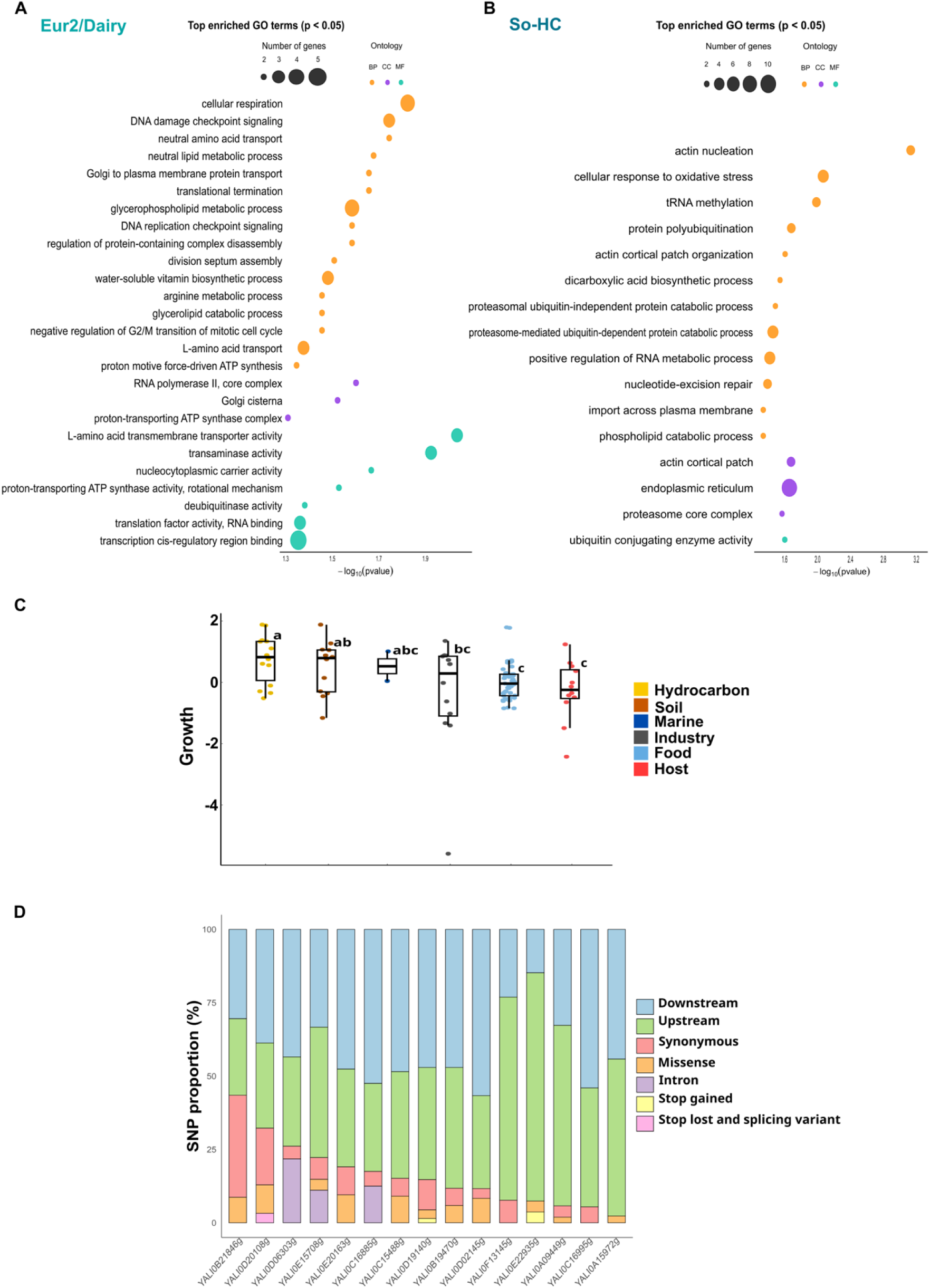
Analysis of population-specific genomic and genetic adaptations. **(A)** Gene Ontology (GO) enrichment analysis of highly differentiated genes (highest mean F_st_ genes) in Eur2/Dairy. **(B)** Gene Ontology (GO) enrichment analysis of highly differentiated genes (highest mean F_st_ genes) in So-HC. Bubble size indicates the number of genes annotated with each enriched GO term; colour indicates GO category: BP (biological process), CC (cellular component), MF (molecular function). **(C)** Growth in acetate as the sole carbon source, normalized by z-score. Strains are grouped and coloured according to isolation source and groups significantly different are classified with letters a,b or c (dunn test). **(D)** Effect of the SNPs located in acetate genes in the top 20% most differentiated genes (F_st_). Gene names are located in the x-axis. Blue (upstream regions) and green (downstream regions) represent regulatory regions.

To illustrate genotype–phenotype links in relevant traits, we focused on growth on acetate as sole carbon source. This trait is of both ecological and biotechnological significance: acetate functions not only as a key environmental metabolite but also as an attractive, low-cost feedstock for the sustainable production of valuable biochemicals. Growth on acetate displays a clear ecological and population-level distribution pattern. Strains isolated from hydrocarbon-associated environments and soils exhibit enhanced growth on acetate compared to those from food-related sources or host-associated niches (Fig. 3C, Kruskal-Wallis test, p = 0.02), consistent with their association to the So-HC group.

To identify genetic variants associated with acetate utilization, we applied two complementary approaches. First, we contrasted the genomes of the 25 best and 25 poorest acetate growers, focusing on the most strongly differentiated genes based on F_st_ values (top 2%). These genes were enriched in functions related to the endoplasmic reticulum (Gene Ontology enrichment analysis, adjusted p = 0.09), including stress response pathways known to contribute to acetate tolerance in yeasts, such as the unfolded protein response, oxidative stress mitigation, and membrane remodeling (Fig. S17). While one differentiated gene was directly linked to acetyl-CoA metabolism (a homologue of IDH2), most genes appeared involved in indirect responses to acetate toxicity. A PCA of haplotypes for these highly differentiated genes revealed a gradient partially aligned with acetate growth performance (Fig. S17). This genetic separation largely mirrored population structure, complicating the disentanglement of evolutionary history versus selection. Nonetheless, some enriched functions differed from those identified in the So-HC group (Fig. 1A), supporting the informative power of our approach.

Second, independently from the genome-wide analysis, we focused on a curated list of 58 genes with known roles in acetate metabolism (Table S5). Applying a more relaxed threshold (top 20% of F_st_ values) within this predefined gene set, we selected 15 divergent genes—including homologues of JEN1, IDH2, and MLS1—for further exploration. The vast majority of associated SNPs in these genes (87%) mapped to non-coding regions (Fig. 3D), suggesting that regulatory changes, rather than alterations to protein sequence, may underlie differential performance.

Altogether, these results exemplify how our dataset enables the dissection of complex traits shaped by both ecology and evolution, and offer a framework for rational molecular bioprospecting based on substrate-specific adaptation.

## Discussion

By integrating population genomics with high-throughput phenotyping, we provide a comprehensive portrait of the natural diversity of *Y. lipolytica*, revealing deep ecological structuring and adaptations to anthropogenic environments.

The population structure of *Y. lipolytica* reflects a complex evolutionary history, with a contrast between genetically diverse, admixed populations and well-defined, specialized lineages. The latter, such as Eur2/Dairy and So-HC, exhibit reduced nucleotide diversity, distinct gene content, and habitat-specific phenotypes, consistent with recent ecological specialization and possible domestication (Duan et al., 2018; Friedrich et al., 2023; Gallone et al., 2016; Vicente et al., 2024). In contrast, populations like Mosaic and Eur1/Mix, sampled from diverse environments, show high genetic diversity, widespread admixture, and evidence of recombination, suggesting a more dynamic evolutionary trajectory shaped by gene flow and admixture (Cromie et al., 2013; Ropars et al., 2018; Tilakaratna & Bensasson, 2017).

Although *Y. lipolytica* is typically found in a haploid state and shows low mating efficiency and spore viability, traits usually associated with limited sexual reproduction, growing genomic evidence challenges the view that clonal reproduction is the sole evolutionary force in this species (Barth & Gaillardin, 1996; Bigey et al., 2023; Li & Alper, 2020; Madzak, 2021). Previous studies reported historical recombination signals, such as short linkage disequilibrium decay (∼50 bp) and introgression (Bigey et al., 2023). Our analyses expand these findings, revealing widespread admixture and gene flow among phylogenetically distant groups (e.g., Mosaic and Eur1/Mix; Fig. 1A), co-ancestry tracts indicative of meiotic exchange (Fig. S9), and signatures of recombination between distinct lineages. These admixed populations form polyphyletic clusters with shared genomic features, underscoring that phylogenetic relatedness alone does not capture their complex evolutionary histories. Together, these patterns support a reticulate evolutionary model, where infrequent but impactful sexual events generate genomic diversity and enhance adaptability (Peris et al., 2014; Ropars et al., 2018; Verhoeven et al., 2011). This resembles observations in *S. cerevisiae*, where gene flow facilitates adaptation to human-associated environments (Duan et al., 2018), as well as in clinical yeasts like *Candida albicans* and *C. glabrata*, previously thought to be asexual but now known to undergo gene flow (Ropars et al., 2018; Wang et al., 2024). Notably, strains isolated from human or animal hosts in our dataset frequently exhibit admixture (11 out of 16), suggesting that host-associated environments may facilitate lineage contact and genetic recombination. Further sampling will be to determine the full extent of this pattern and to identify its ecological drivers. Whether this genetic diversity and ecological versatility represents a stable long-term strategy or a transient state prior to specialization remains to be confirmed through temporal sampling, ecological context, and experimental evolution.

In contrast to admixed populations, structured lineages (Eur2/Dairy, IndNA, So-HC and the W29 clade) show clear signs of environmental adaptation. Nucleotide diversity in these lineages is 10 to 100 times lower than in other *Saccharomycotina* yeasts, such as *S. cerevisiae* (π ≈ 0.003) or *Lachancea thermotolerans* (π ≈ 0.009) (Peter et al., 2018; Vicente et al., 2024), supporting the action of strong ecological filtering or historical bottlenecks. The W29 clade, So-HC, and Eur2/Dairy show high degree of clonality, similar to what has been observed in other fungal species, such as in cheese-adapted *Penicillium* populations (Dumas et al., 2020; Ropars et al., 2020).

Eur2/Dairy strains, isolated from cheese and dairy environments, are enriched in phenotypes advantageous in such contexts, including enhanced lactate utilization and salt tolerance (Mansour et al., 2008; Tunaydın et al., 2024; Viljoen, 2001). They also display reduced lipolytic and proteolytic activity, activities desirable in cheese production to preserve flavor and texture. These traits mirror domestication signatures observed in other fungi (Bennetot et al., 2023) and suggest that *Y. lipolytica* has undergone indirect human-driven selection in this context. Additionally, the most differentiated genes were enriched for amino acid transport functions, aligning with the well-established reliance of *Y. lipolytica* on amino acids as a primary nutritional source in dairy environments (Mansour et al., 2008).

The So-HC lineage, enriched in hydrocarbon-associated strains, shows enhanced growth on hexadecane, ethanol, and acetate, elevated lipase activity, and increased copy number of iron acquisition genes, traits likely beneficial in lipid-rich, iron-limited environments (Salam, 2023). Their close phylogenetic relationship to soil isolates suggests possible transitions between natural and polluted habitats, highlighting the species’ ecological plasticity. Moreover, the most divergent genes and So-HC-specific CNV gains were enriched in oxidative stress response pathways, congruent with the fact that lipid consumption generates reactive oxygen species (ROS), necessitating enhanced stress mitigation mechanisms (Xu et al., 2017).

Together, these findings suggest that *Y. lipolytica* evolves through dual strategies: some lineages undergo genome stabilization and specialization, while others retain reticulate signatures that may facilitate the acquisition of adaptive traits. This evolutionary flexibility may underlie the species’ success across diverse ecosystems, from marine environments and forest soils to dairy products and hydrocarbon wastes.

Beyond evolutionary insight, our dataset offers a robust platform for trait-based bioprospecting and improved application of underutilised strains. The joint genomic and phenotypic landscape enables the exploration of genetic determinants underlying traits of both ecological and industrial relevance. For example, we identified candidate genes and variants associated with enhanced acetate utilization, a substrate of industrial relevance in the biotechnology of this species due to its applicability in lipid and chemical production from low-cost wastes (Chen et al., 2021; Narisetty et al., 2022). These include *S. cerevisiae* transporters orthologs involved in acetate tolerance (*JEN1*, *MCH2*) and known *Y. lipolytica* genes in acetate metabolism such as *IDH2* and *MLS1* (Fu et al., 2024; McCammon, 1996; Salas-Navarrete et al., 2023). Interestingly, most variants associated with the studied acetate-related genes fall within putative regulatory regions. This supports the idea that adaptive evolution often targets gene expression rather than protein sequence, especially for metabolic traits (Fraser et al., 2010; Nourmohammad et al., 2017; Wray, 2007). Regulatory regions may thus offer valuable targets for strain optimization without altering core enzymatic functions. These insights may guide strain selection and inform genomic engineering programs for sustainable biotechnology.

Finally, our results emphasize the importance of considering natural intraspecific diversity when making species-wide inferences. The commonly used reference strain W29 belongs to a distinct, phylogenetically isolated clade, underscoring the limitations of generalizing from a single genetic background.

By delineating the genomic and phenotypic landscape *of Y. lipolytica*, this study advances our understanding of yeast biodiversity and evolution beyond classical fermentation models. In this light, *Y. lipolytica* emerges as a compelling model for studying how ecological interactions, gene flow, and niche specialization shape genome variation and adaptive potential in microbial eukaryotes. Finally, we highlight the untapped potential of natural diversity for bioprospecting, strain engineering, and exploring non-conventional yeasts in biotechnology.

## Limitations of the study

Despite providing a comprehensive view of the population genomics and eco-evolutionary dynamics of *Y. lipolytica*, our study has several limitations. First, the strain collection analyzed is inherently biased toward anthropogenic environments, particularly dairy and industrial niches, which reflects the current composition of global culture collections. This underrepresentation of wild or natural isolates likely limits our ability to fully capture the species’ ecological range and the evolutionary forces shaping its natural populations. Expanding the bioprospecting of *Y. lipolytica* in diverse and underexplored habitats will be essential to uncover novel genotypes and potentially distinct lineages, allowing a more balanced view of its population structure and adaptive landscape.

Second, while we detect signals of admixture, gene flow, and extensive haplotype sharing in certain populations, our inferences on reticulate evolution and potential sexual reproduction are based on genomic patterns alone. A definitive characterization of the reproductive biology of the species, especially the frequency and functional outcomes of outcrossing events, requires experimental validation. Thus, a thorough investigation of reproductive isolation mechanisms and mating compatibility among lineages is necessary to clarify the role of sexual recombination in shaping the observed population structure.

Additionally, although our integrative approach combining pangenome profiling, CNV analysis, and phenotype-genotype associations uncovered candidate adaptive genes, functional validation of specific variants remains to be addressed. Experimental assays (e.g., gene editing or expression profiling under relevant conditions) will be needed to determine the causal relationships between genomic changes and adaptive phenotypes, particularly for traits like acetate tolerance or niche-specific gene losses.

Finally, environmental metadata for some strains were incomplete or inconsistently reported in culture collections, limiting the resolution of eco-evolutionary inferences. Future efforts integrating standardized metadata, ecological sampling, and functional assays will enhance the interpretation of genotype–environment associations.

## Material and Methods

### A collection of *Y.lipolytica* strains

We assembled a comprehensive collection of 126 *Yarrowia lipolytica* strains sourced from various culture collections and research groups worldwide. This effort aimed to capture the broadest possible representation of the species’ ecological diversity. Our collection spans a wide range of geographical regions (America, Asia, North Africa, and Europe) and ecological contexts (food sources, hosts, industrial and urban environments, marine waters, and soils). Notably, European isolates constitute more than half of the collection.

Of the 126 *Y. lipolytica* strains analyzed, 67 were newly sequenced in this study. For the remaining 59 strains, we retrieved raw short-read sequencing data from the NCBI Sequence Read Archive (SRA) (Table S1). The majority of these sequences were originally published by Bigey et al. (2023) and generated using the HiSeq2000 platform (paired-end, 100 bp reads).

### DNA extraction and whole-genome sequencing

Yeasts were grown on YPD agar plates (10 g/L yeast extract, 20 g/L peptone, 20 g/L glucose, 15 g/L agar) at 25 °C for 24 hours. Biomass was collected and treated overnight at 37 °C with Lyticase (Sigma-Aldrich) to digest the cell walls. Chemical lysis by addition of SDS (final concentration 14%) and proteinase K was performed, with incubation at 60 °C for 1 hour. Lysates (520 µL) were then treated with 100 µL 10% CTAB in 0.7 M NaCl and 100 µL 5 M NaCl and incubated at 65 °C for 10 minutes. DNA was purified by extraction with chloroform:isoamyl alcohol (24:1), precipitated with absolute ethanol, washed with 70% ethanol, and resuspended in elution buffer (10 mM Tris-HCl, 1 mM EDTA, pH 8.0). Samples were treated with RNase (3 µL of 10 mg/mL) to remove RNA contaminants. DNA quality was confirmed by gel electrophoresis and spectrophotometry (NanoDrop™ 2000c, Thermo Fisher Scientific). Sequencing was performed using an Illumina NovaSeq 6000 platform generating paired-end 150 bp reads (Novogene).

### Quality control and variant-calling pipeline

We assessed read quality using FASTQC (v0.11.9) and trimmed low-quality bases (q < 20), adapters, and poly-G sequences with Fastp (v0.20.1) (Andrews, 2010; Chen et al., 2018). After trimming, we determined the mating type and ploidy of each strain by mapping reads to reference mating-type loci (*MATA* from W29 strain and *MATB* from E150 strain) using BWA-MEM (v0.7.17-r1188), counting mapped reads per locus, and classifying strains accordingly. One diploid strain (CBS 6614) was identified and excluded from downstream analyses. Filtered reads were then mapped to the revised genome of strain E150 using BWA-MEM with default settings (Bigey et al., 2023; Li & Durbin, 2009). We evaluated alignment quality with Qualimap (v2.2.2) and discarded strains with low mapping rates (<60%) (García-Alcalde et al., 2012).

Alignment files were sorted and indexed using samtools (v1.6), and duplicates were marked with Picard tools (v2.27.4, http://broadinstitute.github.io/picard/). We then applied a GATK-based variant-calling pipeline (v4.1.9) including HaplotypeCaller, GenomicsDBImport, and GenotypeGVCFs to identify single nucleotide polymorphisms (SNPs) and insertions/deletions (indels) (Van der Auwera & O’Connor, 2020). Variants were filtered to retain those with minimum quality and depth thresholds (--minQ 30--min-meanDP 20), excluding indels and non-biallelic SNPs using vcftools (v0.1.16) (Danecek et al., 2011).

### Population Genomics and genetic diversity

We explored the phylogenetic relationships among 126 *Y. lipolytica* strains using 207,153 filtered biallelic SNPs. The data was converted to PHYLIP format and analyzed with IQ-TREE (v2.0.3) to generate a maximum likelihood phylogeny. Model testing was performed to select the best fit, and the tree was built using the ultrafast bootstrap option and ascertainment bias correction (-st DNA-o “outgroup”-m GTR+F+G4-nt 8-bb 1000) (Nguyen et al., 2015). Phylogenetic trees were visualized using the *ggtree* R package (v3.6.2).

Population structure was initially evaluated with ADMIXTURE (v1.3.0) focusing on a subset of unlinked bialellic SNPs (Alexander, Novembre, & Lange, 2009). Filtering of biallelic SNPs based on linkage disequilibrium (LD) was carried out with plink with parameter --indep-pairwise 50 10 0.2 (v1.90b6.21) (Purcell et al., 2007). We initially tested the number of ancestral populations (K) from 2 to 16. The Puechmaille method was used to select the optimal K, as the cross-validation error approach tended to overestimate K in our dataset. This overestimation likely reflects high levels of gene flow and admixture among populations, which complicate the genetic structure and lead to inflated K values with standard methods (Puechmaille, 2016).

To corroborate the population structure, we used fineSTRUCTURE and Principal Component Analysis (PCA). For fineSTRUCTURE, SNP data were formatted with PLINK and chromosome painting was performed using ChromoPainter V2, assuming a constant recombination rate based on *S. cerevisiae* (0.4 cM/kbp) (Cubillos et al., 2011). fineSTRUCTURE (v4.1.1) was run with parameters-x 100000-y 100000-z 1000 (Lawson et al., 2012). PCA was conducted with SMARTPCA from the eigensoft package (v8.0.0) (Patterson et al., 2006).

Overall, genetic groups were defined based on an integrative approach combining the previous lines of evidence. ADMIXTURE analysis, PCA, and fineSTRUCTURE clustering collectively supported the presence of six genetic groups, including four genetically distinct and two admixed (polyphyletic) groups. Phylogenetic relationships from the maximum likelihood tree and patterns of SNP density identified well-supported clades such as the W29 clade, classified as a distinct genetic group due to its clonality and unique genomic profile.

Genetic diversity parameters were calculated using three complementary approaches to capture different aspects of population structure. First, diversity metrics were estimated only for “pure” strains with ancestry proportions >99% according to ADMIXTURE, excluding admixed (mosaic) strains and clonal groups such as the W29 clade. This approach focused on genetically well-defined groups and resulted in 62 strains: 7 from Eur1/Mix (23.3%), 19 from Eur2/Dairy (47.5%), 12 from IndNA (75%) and 24 from So-HC (80%). Second, metrics were calculated including all strains, to assess diversity within admixed groups and the broader population. Third, admixed groups Eur1/Mix and Mosaic were subdivided into phylogenetically defined clades to evaluate whether their diversity patterns were not artefacts of polyphyly. Nucleotide diversity (π) and fixation index (F_st_) were calculated with the *vcfR* R package (v1.15.0). Tajima’s D was calculated genome-wide for each genetic population using VCFtools, employing a window size equivalent to the entire genome length (20.5 Mb) to obtain population-level estimates.

To further investigate historical gene flow and admixture among genetic groups, we calculated D-statistics (ABBA-BABA test) using Dsuite (Malinsky et al., 2021). This approach tests for excess allele sharing among quartets of populations (P1, P2, P3, and an outgroup) to detect introgression signals. We used LD-filtered SNP dataset of “pure” strains and defined parental groups based on phylogenetic and population structure results. Statistical significance was assessed using Z-scores (>3) and p-values (< 0.05).

### Gene content analysis

Copy number variants (CNVs) and pangenome analysis were utilized to explore gene content variations across our collection. Control-FREEC (v11.6), a tool leveraging read coverage and GC content, was employed to study CNVs (parameters: window = 1000; breakPointThreshold = 0.05; minExpectedGC = 0.35; maxExpectedGC = 0.55; ploidy = 1) (Boeva et al., 2012). Results were visualized with the R package pheatmap (v1.0.12).

Absent genes in the reference strain were identified following a previously described pipeline for the analysis of the pangenome (Gounot et al., 2020). The presence of a low percentage of reads (2%) belonging to the *Cellulosimicrobium* genus was detected, so modifications were incorporated into the pipeline to eliminate non-*Y. lipolytica* sequences. Briefly, quality-filtered reads were aligned to the reference genome of *C. cellulans* with BWA, as described in previous sections, and unmapped reads were extracted using samtools. Then, genomes were assembled with SPAdes (v3.13.1), using automatic detection of k-mer size. Non-reference genetic material was identified by comparison against the reference strain with Blast (v2.16.0) and genes were predicted using SNAP and AUGUSTUS (v2.5.5), with a trained model against reference strain and a pre-existing model, respectively (Korf, 2004; Stanke, Schöffmann, Morgenstern, & Waack, 2006). Low-complexity regions and proteins shorter than 50 amino acids were excluded from the analysis. The remaining proteins were compared against those from the *Cellulosimicrobium* genus to eliminate potential foreign reads using BlastP. Finally, redundant genes were grouped into genetic clusters through a graph-based method using pairwise similarity scores, with a default similarity threshold of 98%, enabling the identification of non-reference material across the dataset.

### High-throughput phenotyping

We phenotyped 105 strains of our collection by evaluating growth and other traits of industrial and ecological interest in 49 conditions encompassing different carbon and nitrogen sources and several environmental stresses. The majority of conditions were evaluated through liquid cultures in microplate growth assays, while a minority of conditions were assessed in solid cultures (Table S3).

Each condition was based on variations of a Standard Medium composed of Yeast Nitrogen Base (YNB 0.67%, BD Difco™, USA) and glucose (1%). Modifications included the substitution of 1% alternative carbon sources, supplementation with specific inhibitors, or changes of environmental variables such as pH or temperature. For experiments testing alternative nitrogen sources, 0.17% YNB without amino acids and ammonium sulfate (Thermo Fisher Scientific, USA) was used, supplemented with the appropriate nitrogen source and glucose (1%). The Standard Medium was further supplemented with 1.5% agar for solid culture assays.

For growth assays in liquid media, yeast strains were pre-cultured in YNB (0.67%) with glucose (0.5%) in 24-well plates at 25 °C for 24 hours. Pre-cultures were then diluted in saline solution (0.9% NaCl) to an optical density at 600 nm (OD_600_) of 0.02. A total of 25 µL of this suspension was inoculated in 48-well plates in triplicate into 475 µL of the corresponding media using a robotic system (Assist Plus – VIALAB system, Integra Bioscience, Switzerland), resulting in an initial cell density of 10⁴ cells/mL. Cell growth at 25 °C (unless temperature was a variable) with orbital shaking at 100 rpm was monitored over 63 hours by measuring OD_600_ using a microplate reader (Varioskan Flash Multimode Reader, Thermo Fisher Scientific, USA). To evaluate pyomelanin production, cells were grown in *Yarrowia lipolytica* Differential growth medium (YLD) and categorized according to brown colour intensity (Akpınar, Uçar, & Yalçın, 2011b)

For solid growth assays, yeast cells were precultured and diluted as before and 10 µL of 10⁴ cells/mL solutions were inoculated into squared Petri dishes (Gosselin^TM^, Thermo Fisher Scientific, USA) in spots and by triplicate. Cells were incubated at 28 °C and pictures were taken after 240 hours of growth. Growth was compared to residual growth in Standard Medium without carbon source and classified into 3 categories: no growth, weak growth or growth. In the case of skimmed milk Agar (Sigma-Aldrich), the absence of a halo was categorized as a lack of protease activity. Colony morphology was evaluated in YPDA (1% yeast extract, 2% peptone, 2% glucose, 1.5% agar) at 25 °C after 48 hours of growth and 3 distinct morphologies were categorised (smooth, intermediate roughness and high roughness).

Lipase activity was measured in 5 different conditions (Standard Medium, Glycerol as carbon source, Urea as nitrogen source, pH 7.0 and C/N ratio = 100 as environmental stresses) at the end time point of cultures by the p-nitrophenyl-butyrate (pNPB) method. Briefly, aliquots of 20 µL were transferred from the cultures at 63 h using the previously mentioned robotic system and transferred to a mix of Tris-HCl 20 mM pH7 buffer and pNPB 1,5 mM. OD_405_ measurements were taken every 2 minutes for 20 minutes, and changes in optical density were used to adjust a linear regression model and quantify the hydrolysis of pNPB. One unit of lipase activity was determined as the amount of enzyme releasing 1 μM of p-nitrophenol per minute under assay condition.

### Phenotypic data analysis

For liquid growth assays, growth data was fitted to the Baranyi model using the R package *biogrowth* (v1.0.4), and key growth parameters such as lag phase, maximum growth rate (µmax), and proliferative efficiency (OD_600_ at various time points) were extracted, eventually choosing proliferative efficiency at 48 hours as our working variable, as proxy for both µmax and proliferative efficiency, and referring to it as growth. The z-score of growth per condition was calculated for each strain and utilized for subsequent data analysis. For conditions testing resistance to different inhibitors and adverse environmental conditions, the growth of each strain was further normalised by the growth in Standard Medium to study the resistance of the strain as a deviation from its standard growth.

Phenotypic distance was calculated as the Euclidean distance among strains using base R (v4.2). For this analysis, we used our non-categorical variables, i.e., the growth in liquid culture in 34 conditions and the lipase activity data. Its correlation with the phylogenetic distance, calculated as the sum of the branch lengths between each pair of strains, was based on Pearson’s correlation. Phylogenetic signal was estimated by determining the Pagel’s λ value using *ape* (v5.7.1) and *phytools* (v2.0.3) R packages as previously described by Ruiz et al. (2023).

To assess phenotypic robustness at the population level, we identified the top 20% of strains per condition based on normalized growth (z-score), and calculated the frequency of these top performers per population, normalizing by group size.

Data manipulation, statistical data analysis and hypothesis testing were performed using base R. PERMANOVA analysis was based on the R vegan package (v2.6.8). Data visualization was carried out with ggplot2 (v3.1.5) and pheatmap (v1.0.12) R packages.

### Genetic basis of population-specific adaptations

To elucidate the genetic basis underlying phenotypic adaptations in the Eur2/Dairy and So-HC populations, we compared each focal population against all other strains to identify the most divergent genes using the fixation index (F_st_). For each analysis, SNPs were extracted using vcftools (v0.1.16) (Danecek et al., 2011), and pairwise F_st_ values were calculated contrasting the focal group against the rest of the dataset. Mean per-gene F_st_ values were computed, and the top 2% of genes with the highest differentiation were selected for downstream analyses. Functional enrichment of these candidate gene sets was performed through Gene Ontology (GO) analysis using the *topGO* (v2.59.0) package in R (Alexa & Rahnenfuhrer, 2010). Additionally, CNV gains unique to these populations were identified and subjected to GO enrichment to further explore population-specific functional signals.

### Genetic basis of acetate performance

To investigate the genetic basis of variation in acetate utilization, we contrasted the genomes of the 25 best and 25 poorest acetate growers, as determined from phenotypic assays, using the same F_st_-based approach described above. Haplotypes of the most highly differentiated genes were reconstructed and analyzed by PCA using the *prcomp* function in R to explore allele distribution patterns.

In parallel, a manually curated list of 58 genes involved in acetate metabolism and acetate-induced stress response (Table S5), based on prior literature in *S. cerevisiae* and other yeasts, was compiled. For genes ranking in the top 20% of F_st_ values, functional annotation of SNPs was performed using SnpEff (v5.2f), allowing assessment of both variant effect type and predicted impact (Cingolani et al., 2012).

## Resource availability

Sequences used and generated in this study can be found in the National Center for Biotechnology Information database under BioProject accession numbers PRJEB42834 and PRJNA1280552 and SRA accession numbers SRR21430003, SRR6475361, SRR6475466, SRR6475467, SRR6475649, SRR6820826 and SRR6820828. Pipelines of analysis and pertaining scripts are available at https://github.com/sergigea689/Yarrowialipolytica_genomics.

## Supporting information

Izquierdo-Gea et al. 2025 - supplementary tables

Izquierdo-Gea et al. 2025 - supplementary figures

## Acknowledgements

Recent research in I.B.’s laboratory has been supported by the Spanish State Research Agency/Ministry of Science and Innovation (https://doi.org/10.13039/501100011033) through grants PID2019-105834GA-I00 (Wineteractions) and PID2022-138343NB-I00 (INDUSYNCON), with co-funding by the European Regional Development Fund (ERDF/EU). S.I.-G. acknowledges funding from the Spanish Ministry of Science, Innovation and Universities through a predoctoral grant (FPU21/06830), and thanks the European Molecular Biology Organization (EMBO) for a Scientific Exchange Grant that supported the collaborations leading to this study. F.C. received funding from ANID–Millennium Science Initiative Program (ICN17_022). J.B. acknowledges funding from the European Union’s Horizon Europe Research and Innovation Program through the CAPTUS (HE 101118265) and FUELPHORIA (HE 101118286) projects. We thank Cécile Neuveglise and Santiago Ruiz-Moyano for kindly providing strains used in this study. The funders had no role in study design, data collection and analysis, decision to publish, or preparation of the manuscript.

## Declaration of interests

The authors declare no competing interests

## Notes

### Competing Interest Statement

The authors have declared no competing interest.

