## Supplementary material for "Contrasting evolutionary forces of specialization and admixture underlie the genomic and phenotypic diversity of *Yarrowia lipolytica*": Izquierdo-Gea et al. 2025 - supplementary figures

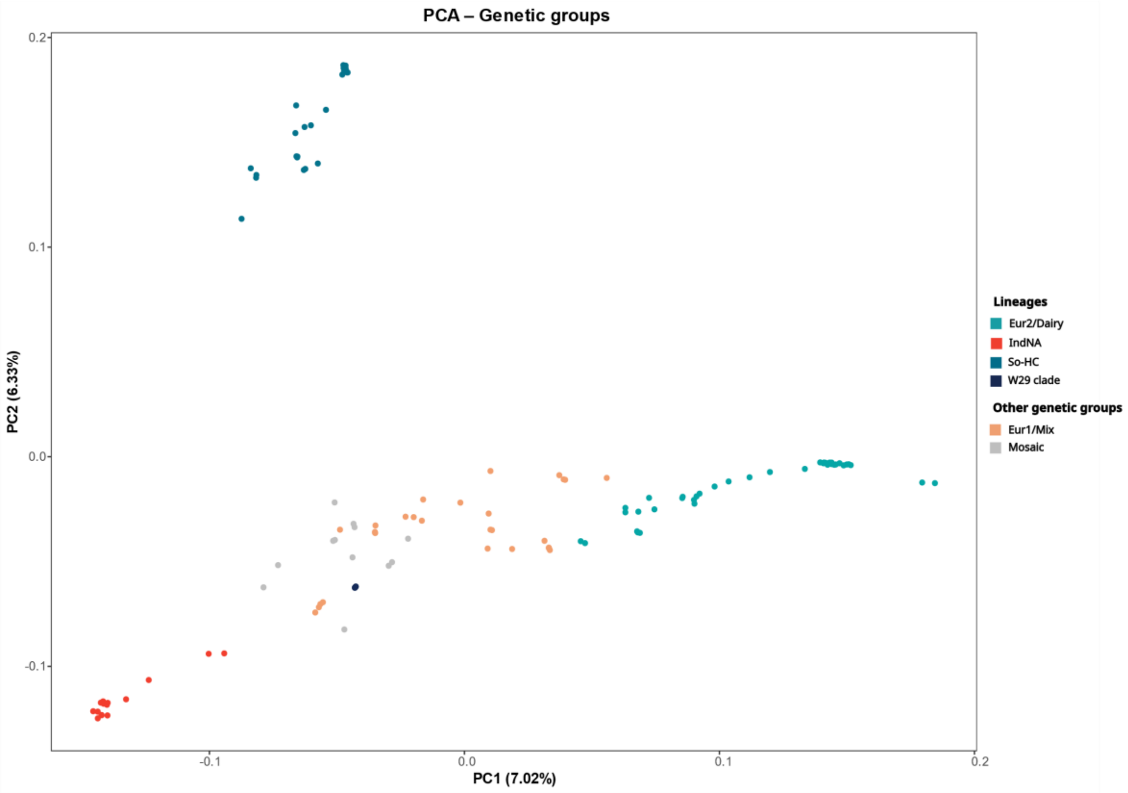


**Figure S1.** Principal component analysis (PCA) of high-quality biallelic LD-pruned SNP data.Principal components 1 and 2 (PC1 and PC2) are shown, with the percentage of variance explained indicated in parentheses. Colors represent genetic groups, either well-defined and structured (Lineages) or less defined, admixed groups (Other genetic groups).

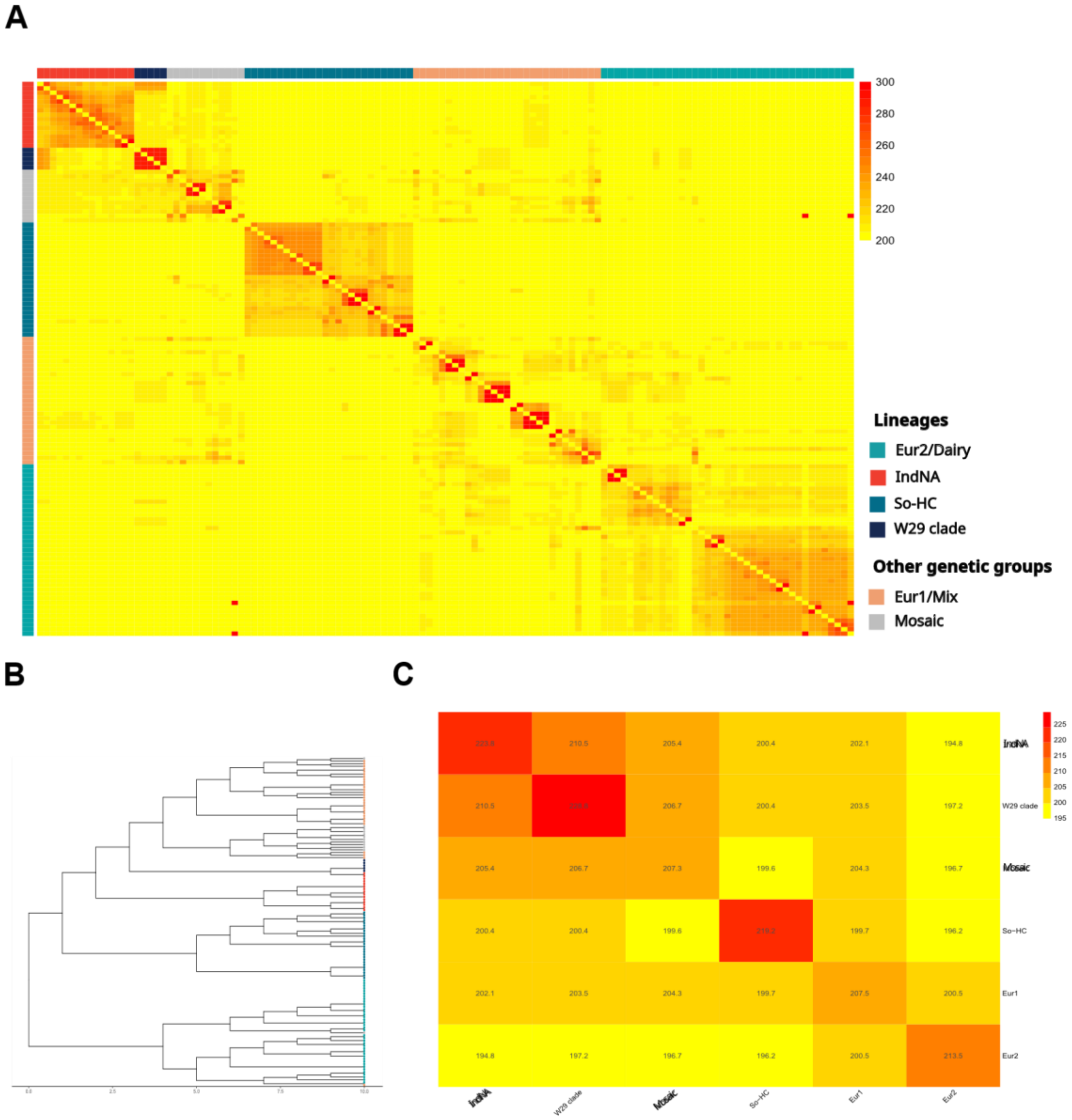


**Figure S2.** Co-ancestry analysis (haplotype similarity) based on fineStructure. **(A)** Co-ancestry heatmap displayed according to tree order and labeled with genetic groups on both the x- and y-axes. Cells represent co-ancestry between each pair of strains. Darker squares indicate co-ancestry clusters. **(B)** Co-ancestry dendrogram, showing samples grouping based on haplotype conservation. Tip points are colored according to the genetic groups labeled in panel A. **(C)** Mean co-ancestry heatmap showing pairwise values among the six genetic groups. Co-ancestry values are shown within the cells.

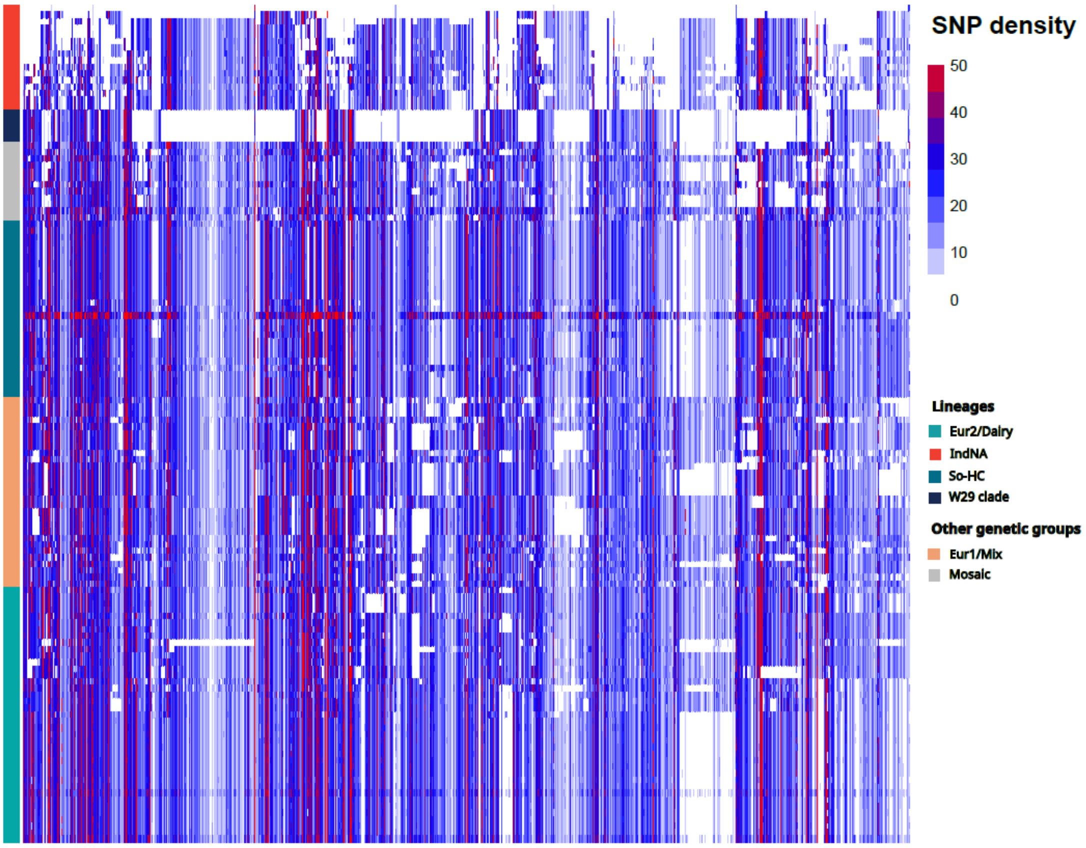


**Figure S3.** SNP density map. Samples are shown on the y-axis, ordered by phylogenetic position and labeled according to genetic group. Genomic windows are displayed on the x-axis in chromosomal order. SNPs were counted in windows of 2,000 bp.

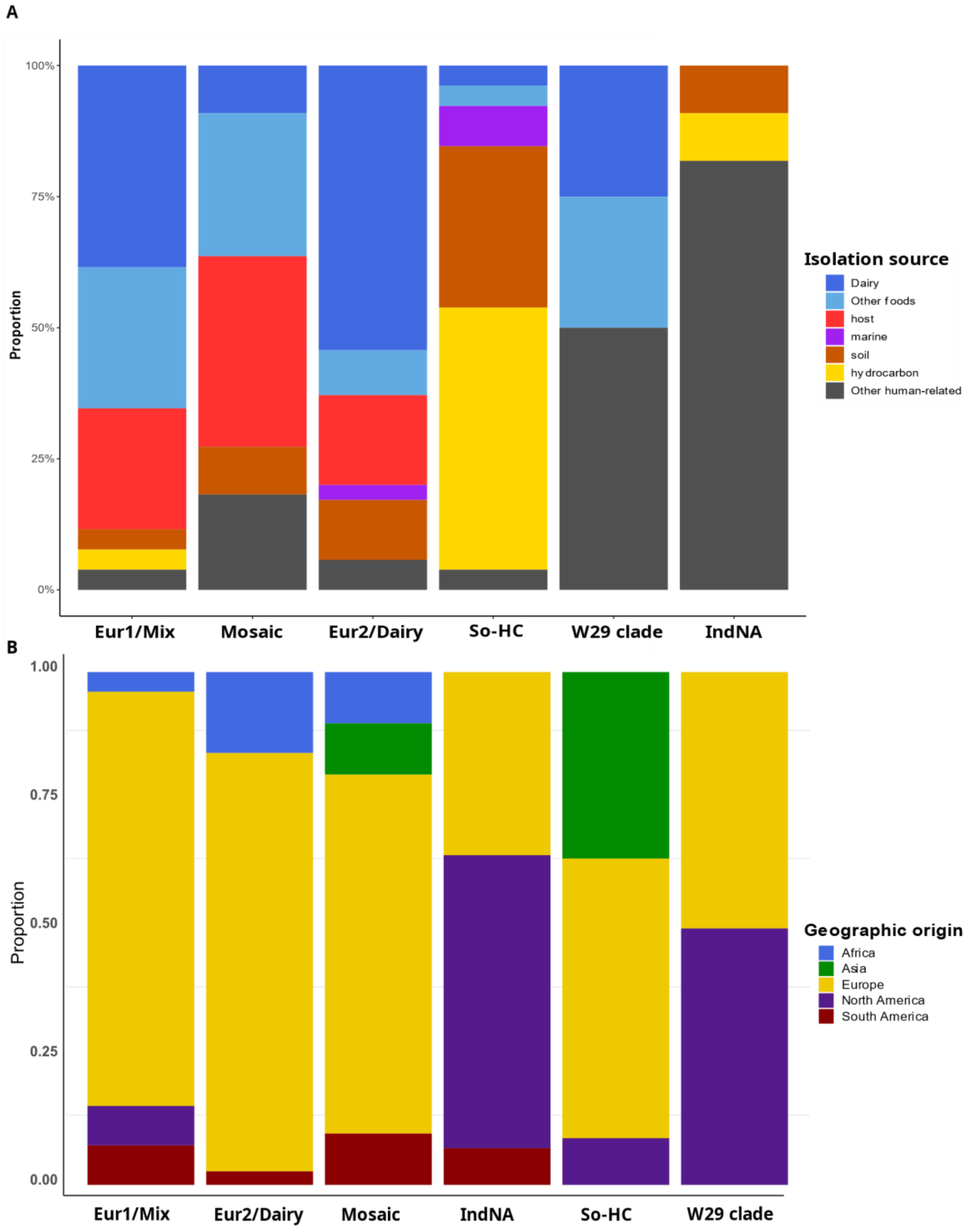


**Figure S4.** Proportion of isolation source **(A)** and geographic origin **(B)** per genetic group.

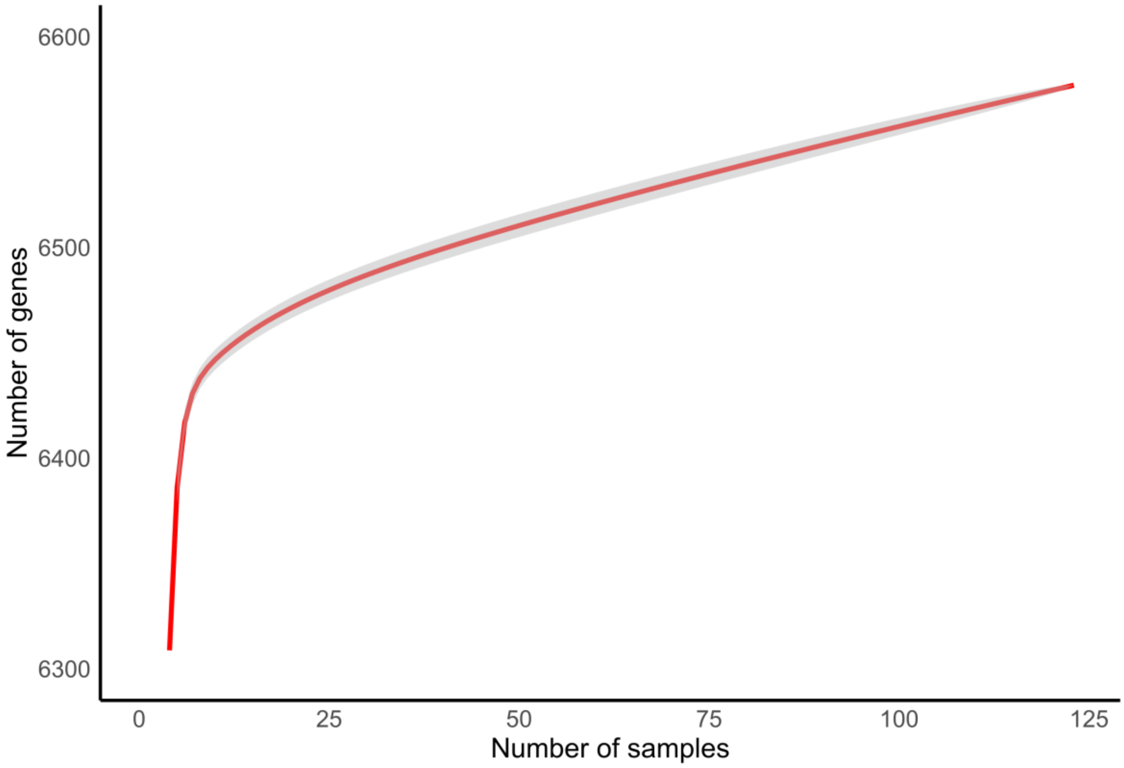


**Figure S5.** Pangenome curve generated from accessory genome analysis of 125 *Y. lipolytica* strains. The curve shows the cumulative number of gene families as a function of the number of genomes analyzed.

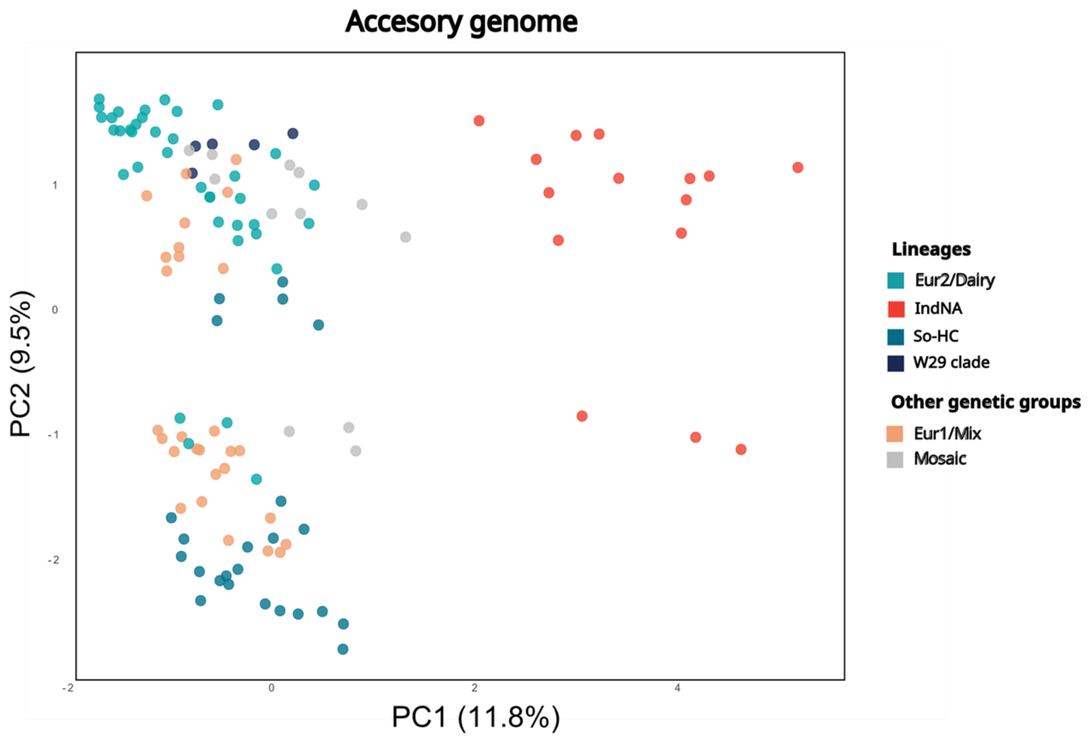


**Figure S6.** PCA analysis of accessory gene content, based on gene losses and non-basal core genes. Samples are coloured according to genetic groups and PC1 and PC2 are displayed, with the percentage of variance explained in parentheses.

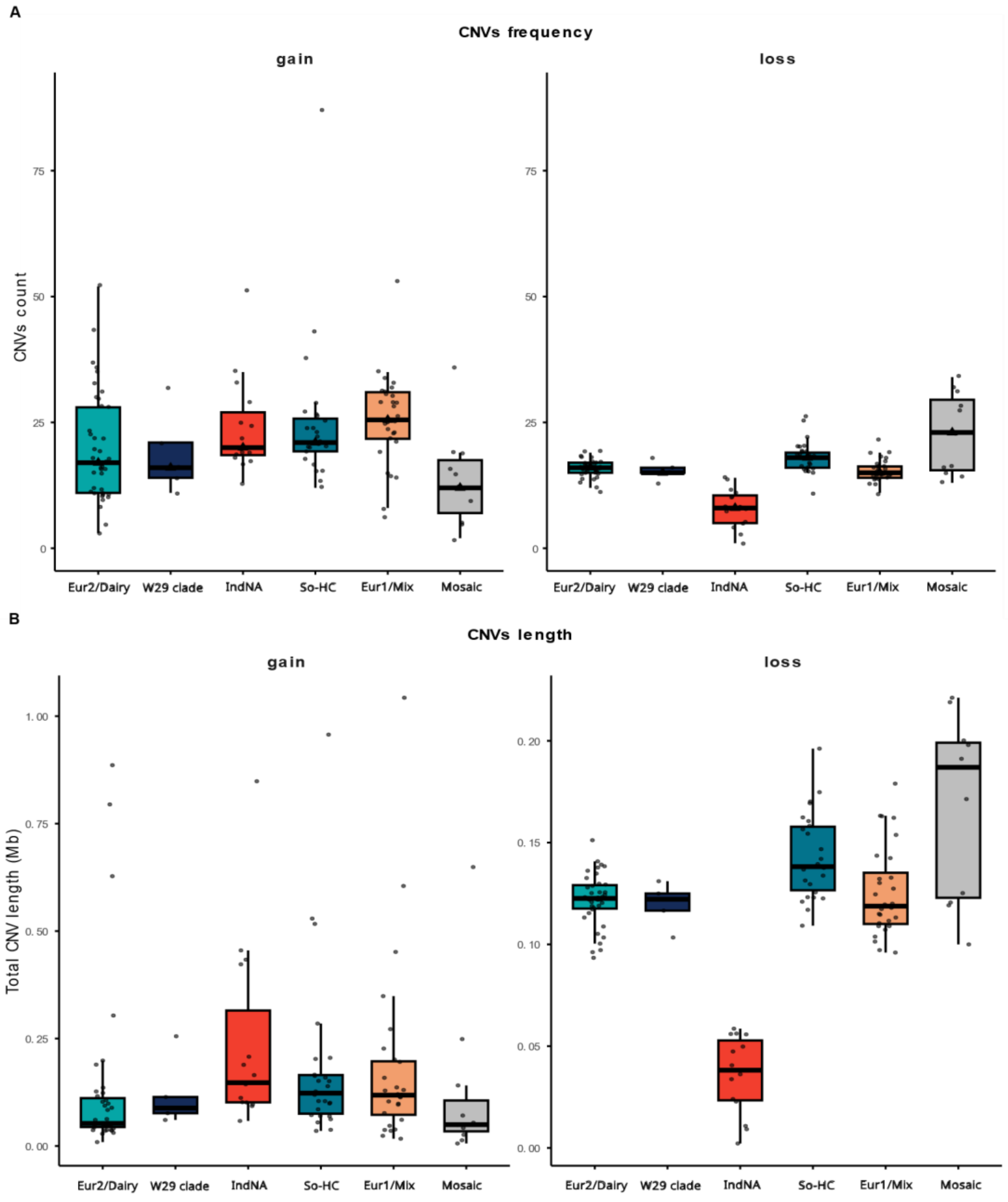


**Figure S7.** Copy number variants (CNVs) analysis per genetic group, corresponding to the colour of each boxplot. **(A)**. CNVs counts per genetic group: gene gains in the left panel and gene losses in the right panel. **(B)**. Total genomic length in millions of base pairs affected by CNVs per genetic group: gene gains in the left panel and gene losses in the right panel.

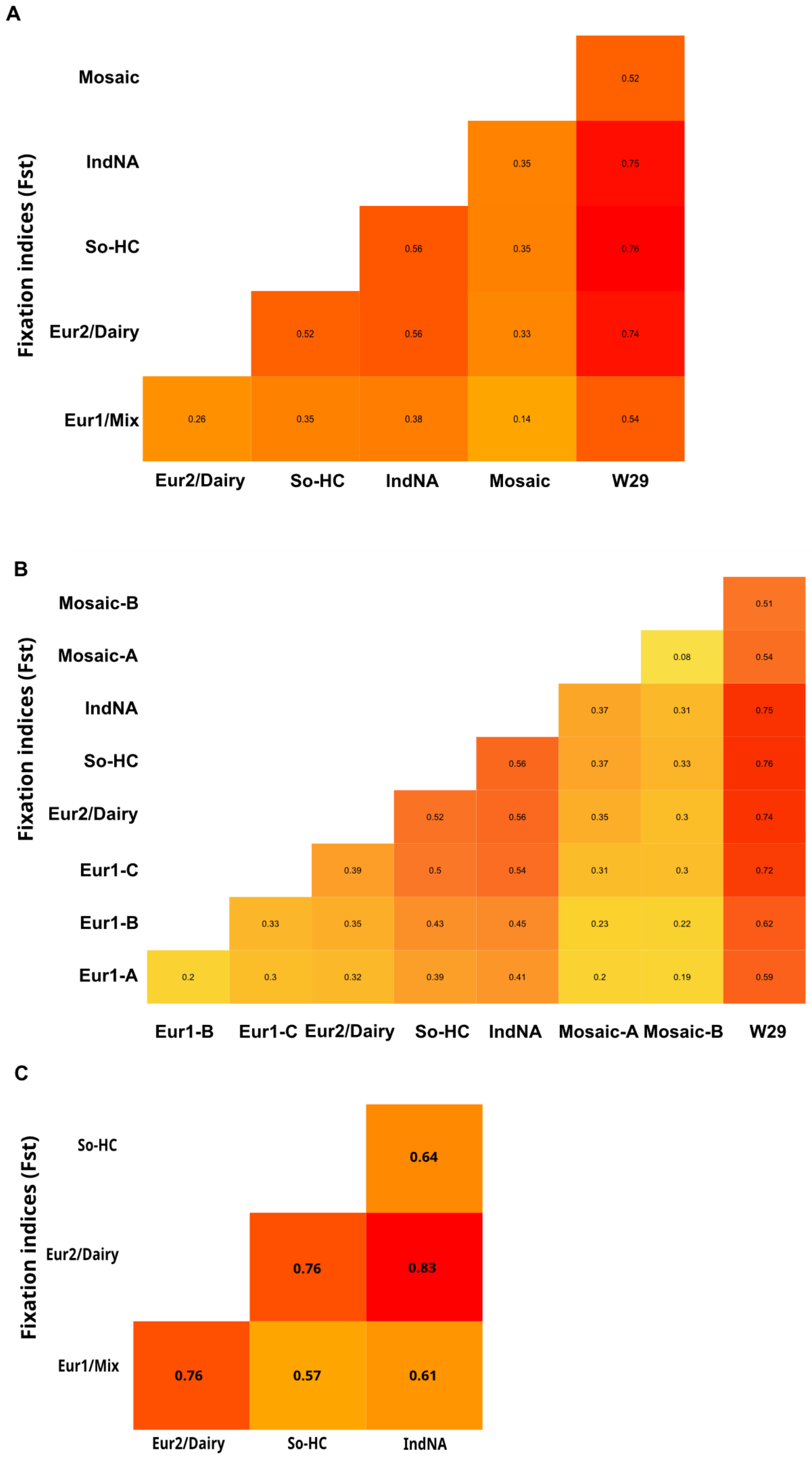


**Figure S8.** Fixation index ($F_{st}$) analysis, showing pairwise differences in allele frequency. Values are shown within the cell. **(A)** Analysis based on every strain of each genetic group. **(B)** Analysis considerating every phylogenetic clade within the admixed groups. **(C)** Analysis including only pure strains (ancestry proportions >99% according to ADMIXTURE), excluding the W29 clade due to its smal size and high clonality

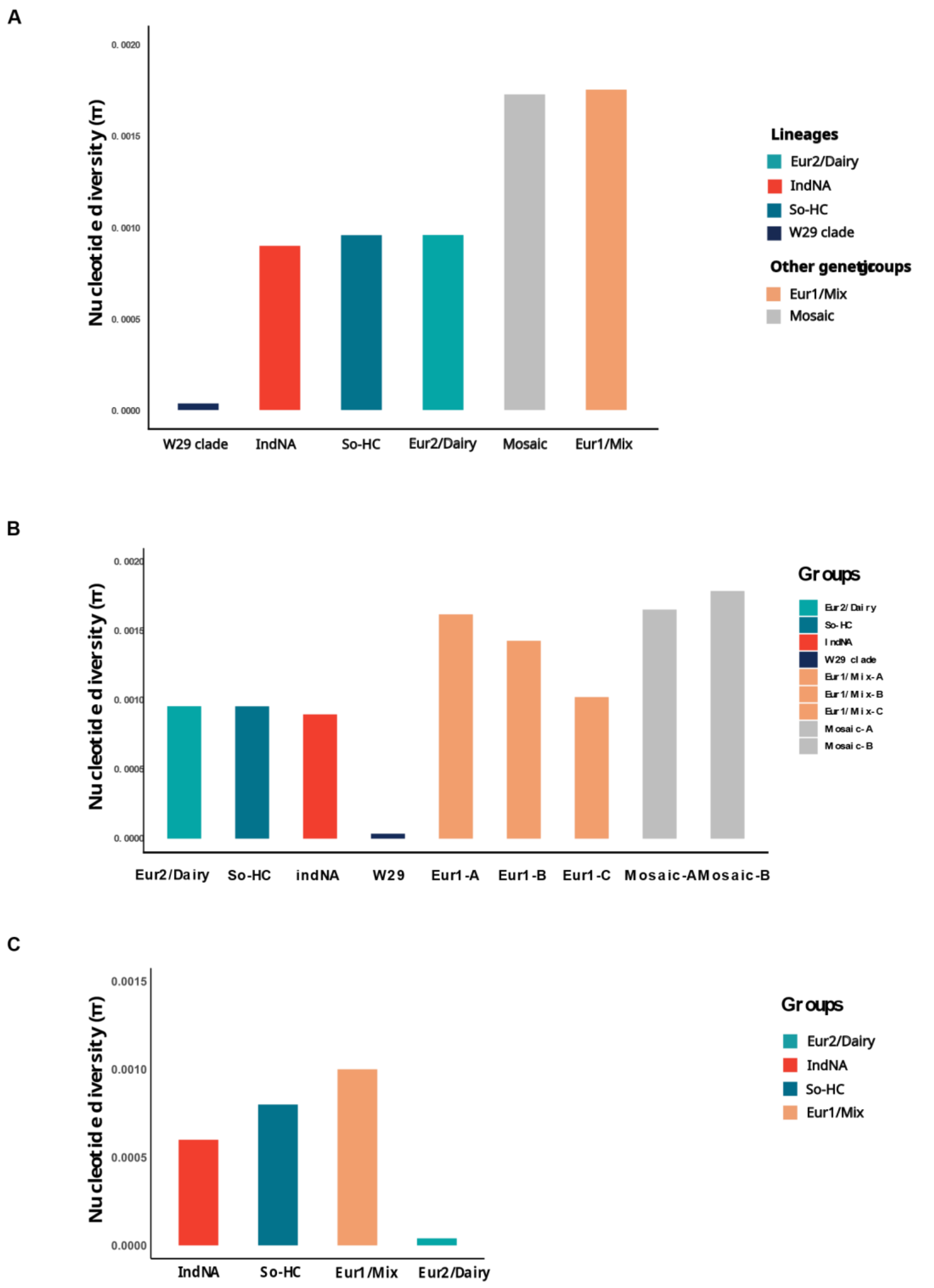


**Figure S9.** Nucleotide divergence (π) analysis, showing intrapopulation nucleotide differences. Bars are coloured according to genetic group. **(A)** Analysis based on every strain of each genetic group. **(B)** Analysis considerating every phylogenetic clade inside the admixed groups. **(C)** Analysis including only pure strains (ancestry proportions >99% according to ADMIXTURE), excluding the W29 clade due to its smal size and high clonality

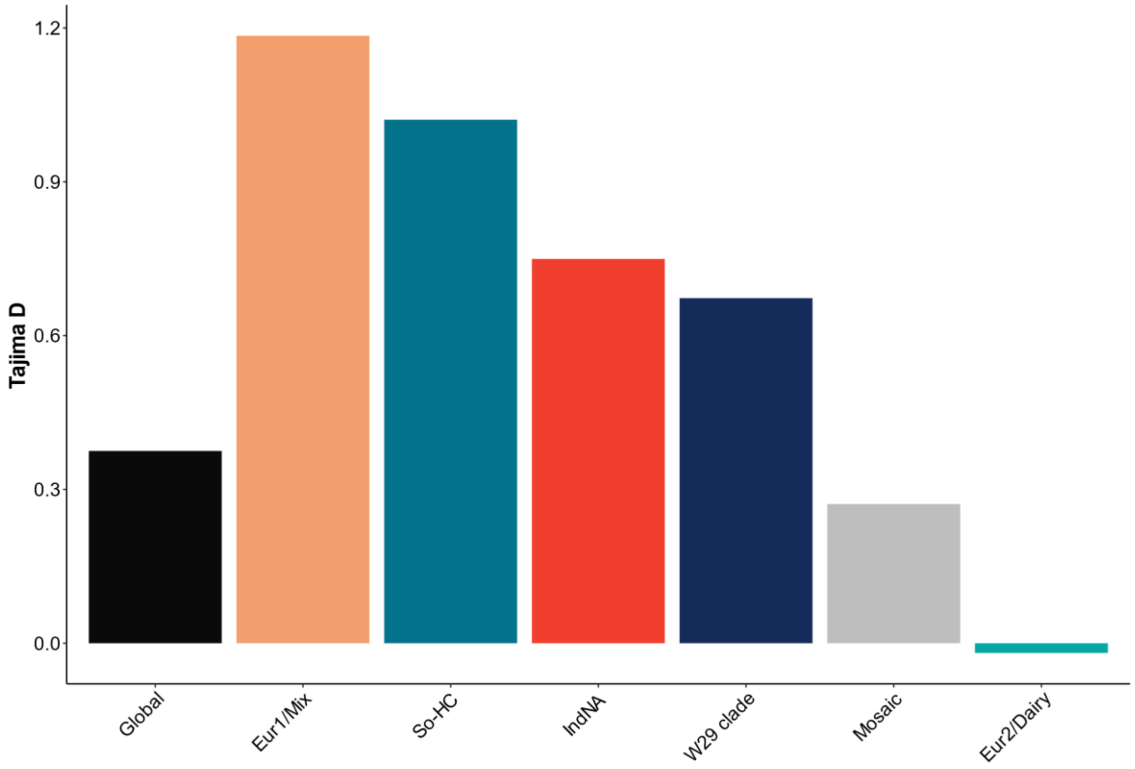


**Figure S10.** Tajima`s D for each genetic group and for the global sample.

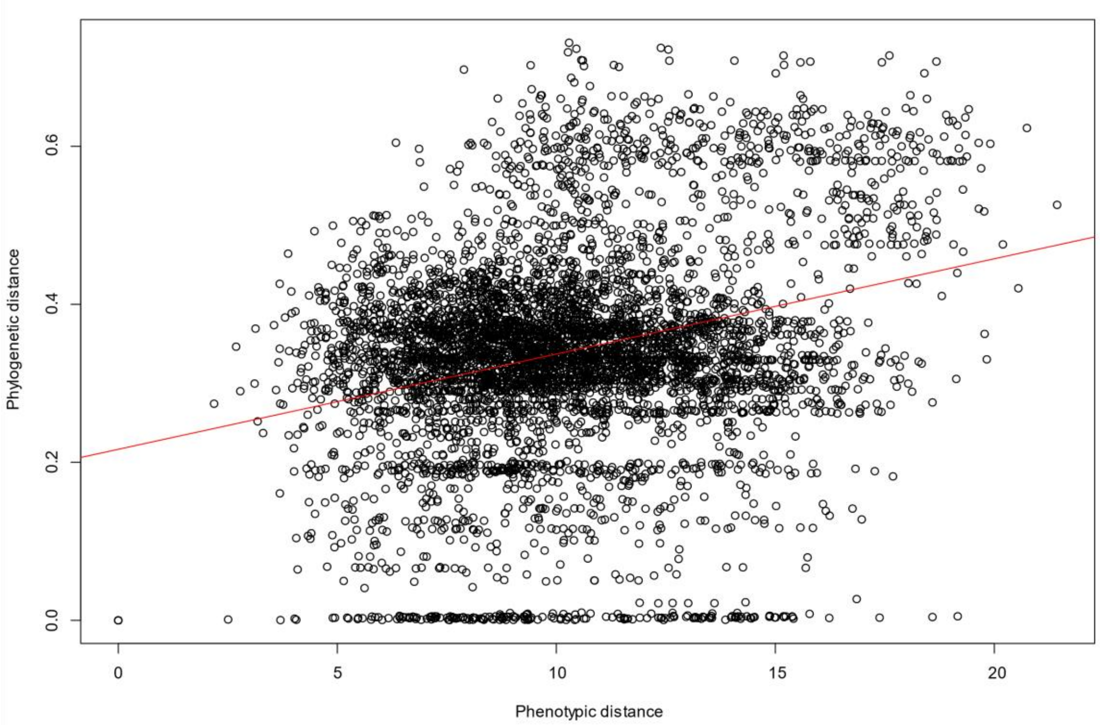


**Figure S11.** Correlation between phylogenetic distance, based on the biallelic SNP phylogeny, and phenotypic distance, based on quantitative growth measurements, across 126 Y. lipolytica strains.The correlation coefficient is r = 0.30, with a significance level of $p = 2.2 \times 10^{-16}$.

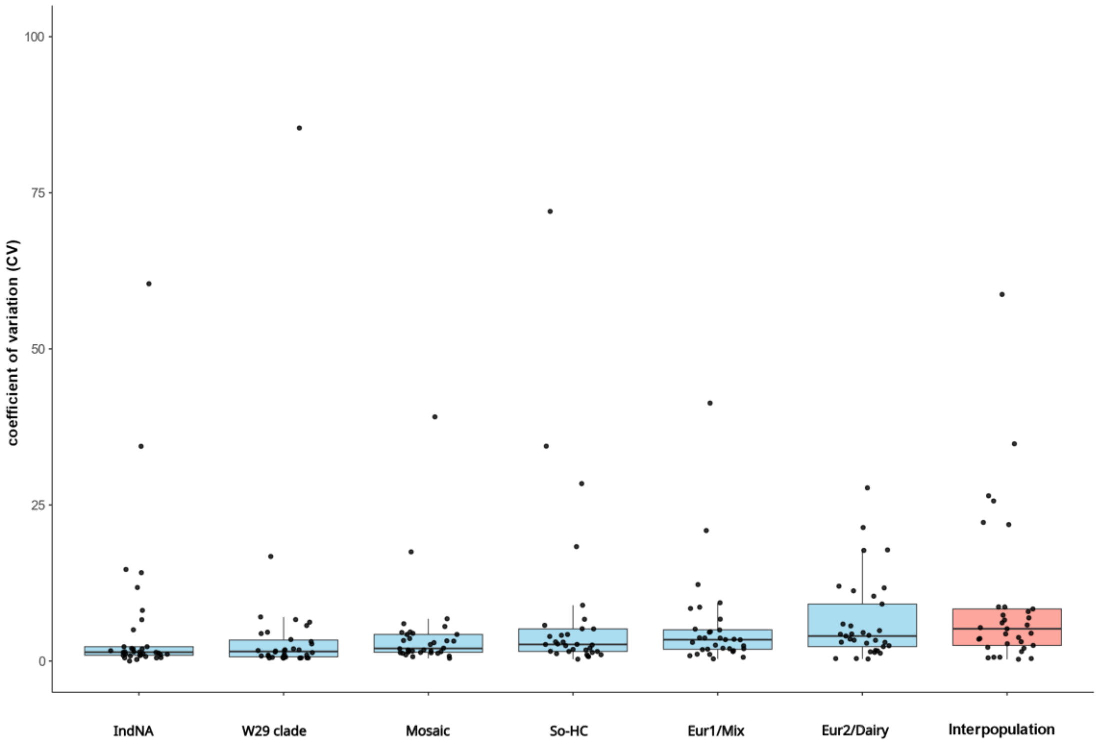


**Figure S12.** Coefficient of variation (CV) of phenotypic measurements within each genetic group (blue) and among genetic groups (pink). Boxplots represent the distribution of CV values across all phenotypic conditions.

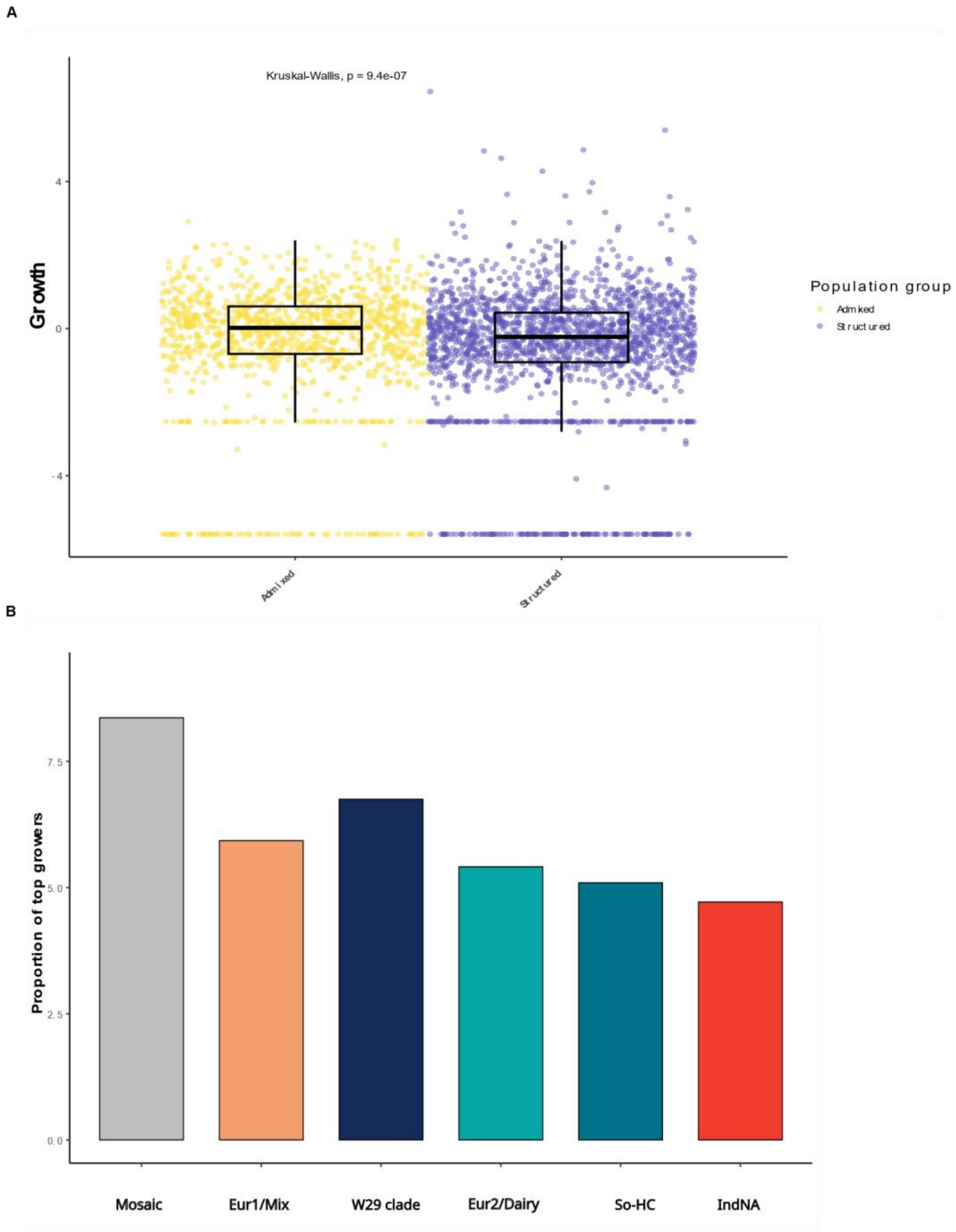


**Figure S13.** Phenotypic data comparing admixed groups and well-defined, structured lineages. **(A)** z-score of growth in every condition of quantitative growth. Kruskal Wallis test showing the difference among groups and corresponding p-value are shown. **(B)** Proportion of strains among 1% of top growers normalised by genetic group sample size.

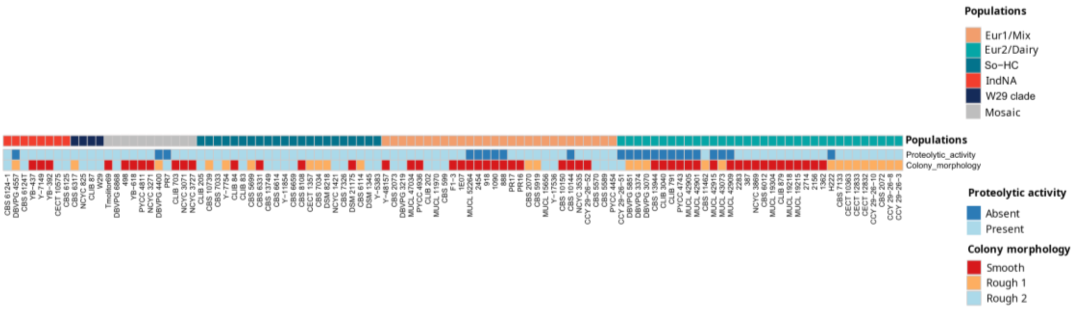


**Figure S14.** Heatmap representation of protease activity and colony morphology, with samples ordered phylogenetically and labeled according to genetic group (populations).

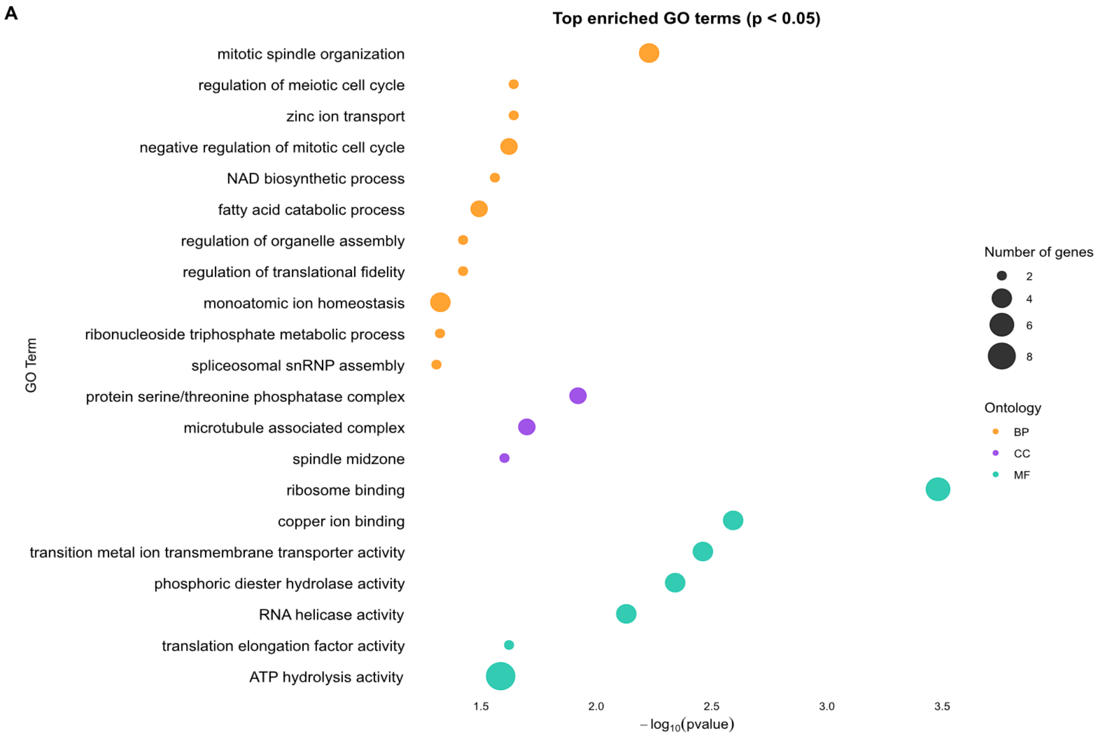


**Figure S15.** Gene Ontology (GO) enrichment analysis of unique CNVs gains of the Eur2/Dairy. Bubble size indicates the number of genes annotated with each enriched GO term; colour indicates GO category: BP (biological process), CC (cellular component), MF (molecular function).

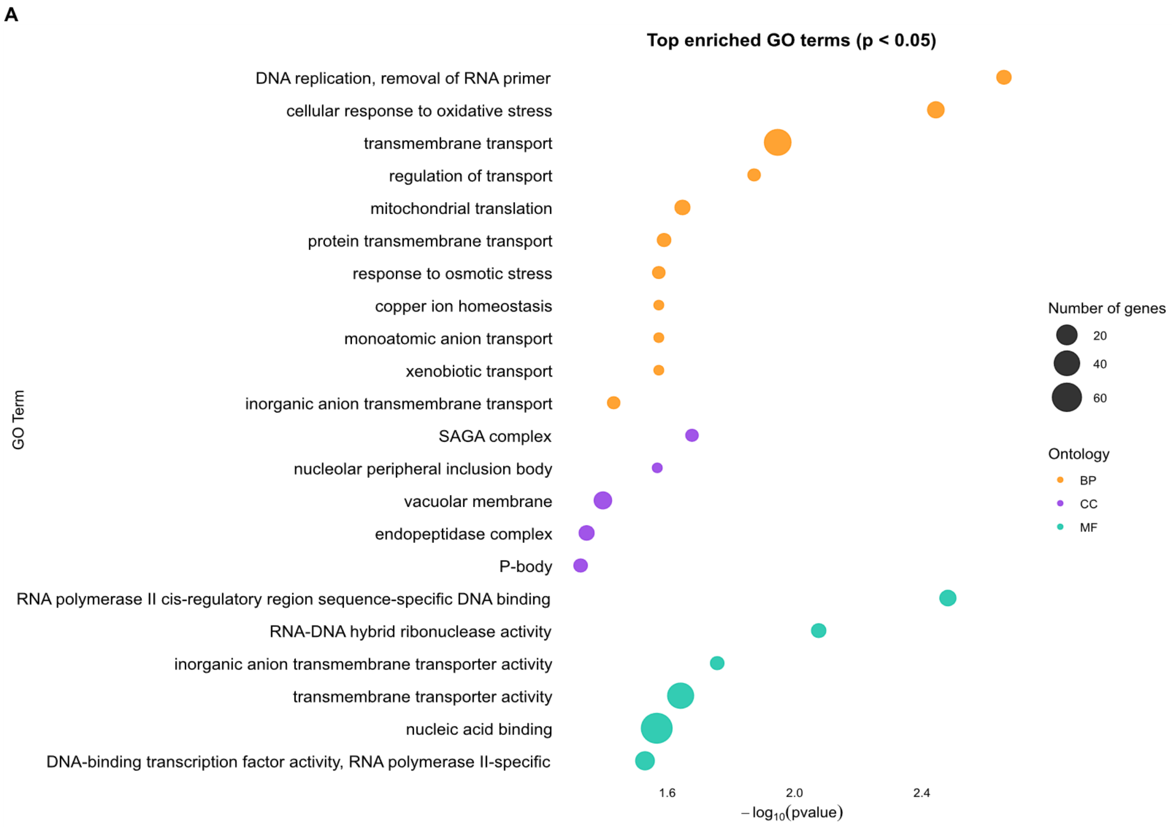


**Figure S16.** Gene Ontology (GO) enrichment analysis of of unique CNVs gains of the So-HC. Bubble size indicates the number of genes annotated with each enriched GO term; colour indicates GO category: BP (biological process), CC (cellular component), MF (molecular function).

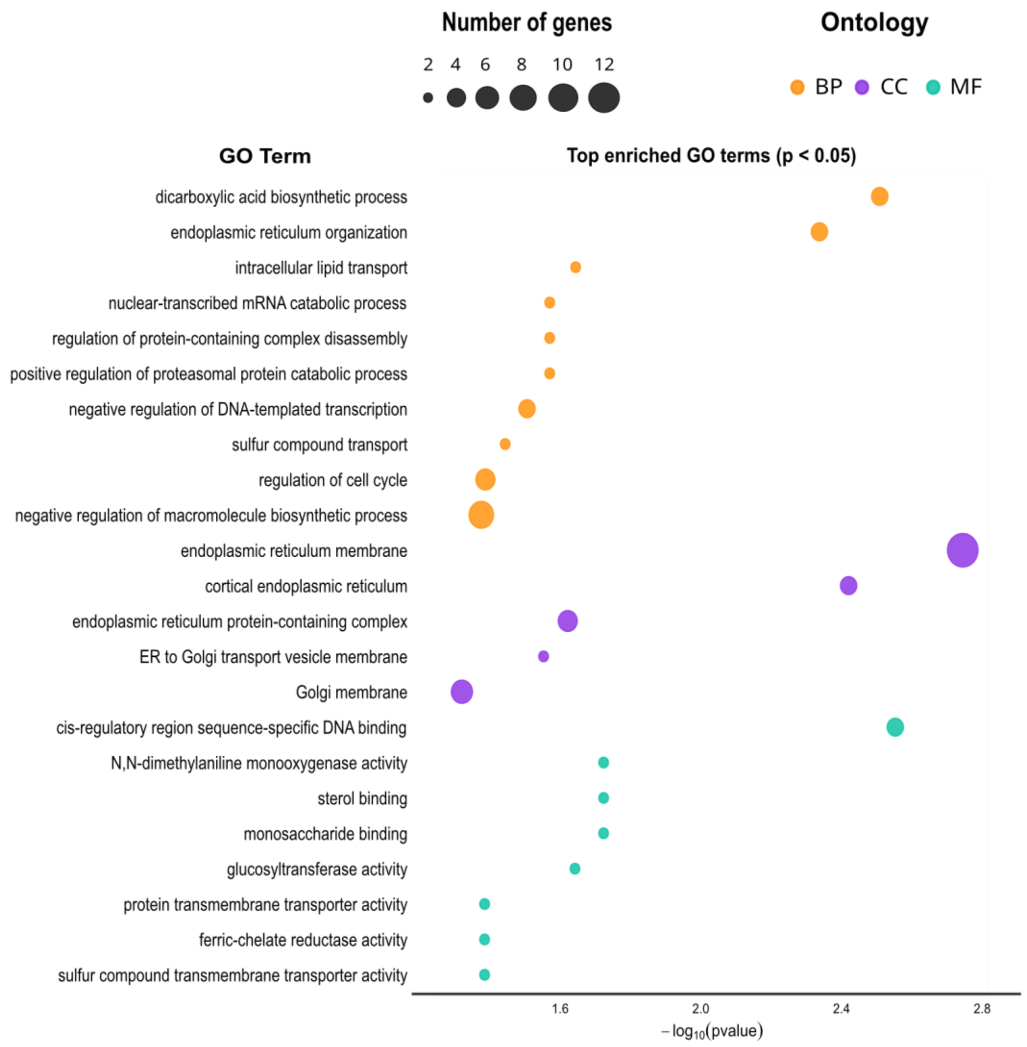


**Figure S19.** Gene Ontology (GO) enrichment analysis of highly differentiated genes among best and worst acetate growers (highest mean $F_{st}$ genes). Bubble size indicates the number of genes annotated with each enriched GO term; colour indicates GO category: BP (biological process), CC (cellular component), MF (molecular function). BP corresponds

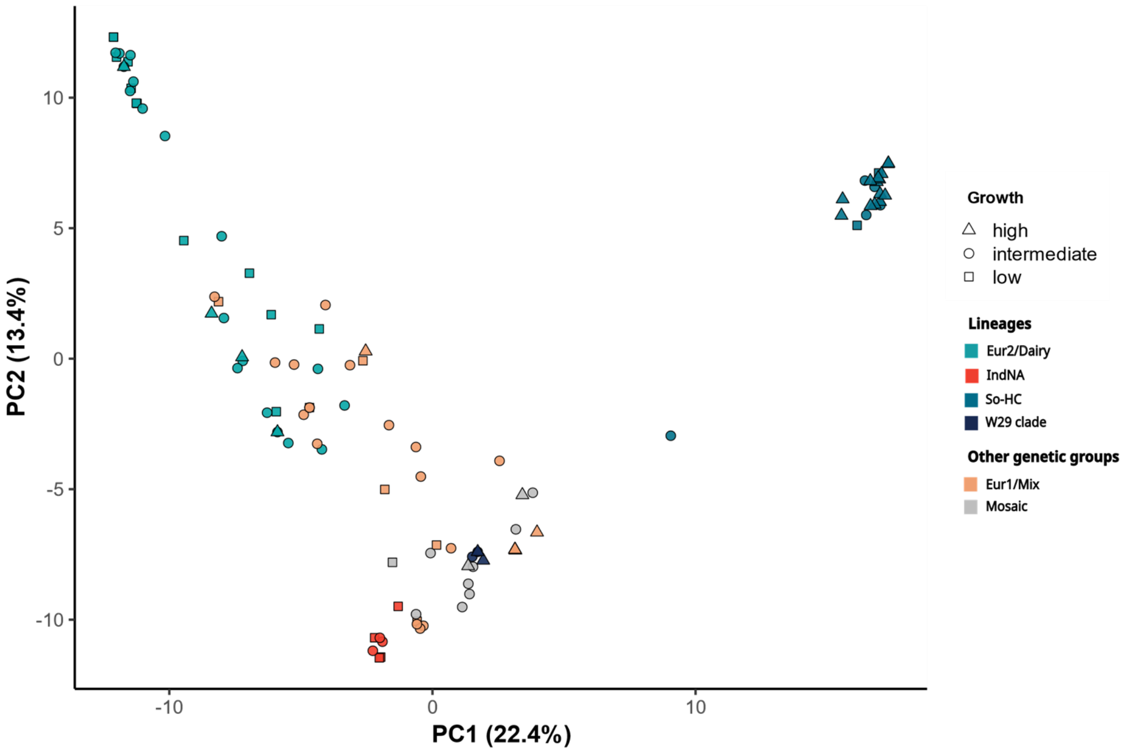


**Figure S18.** PCA of the most abundant haplotypes from the most differentiated genes between the best and worst acetate growers. Samples are colored according to genetic group, and shapes indicate growth performance in medium containing acetate as the sole carbon source.

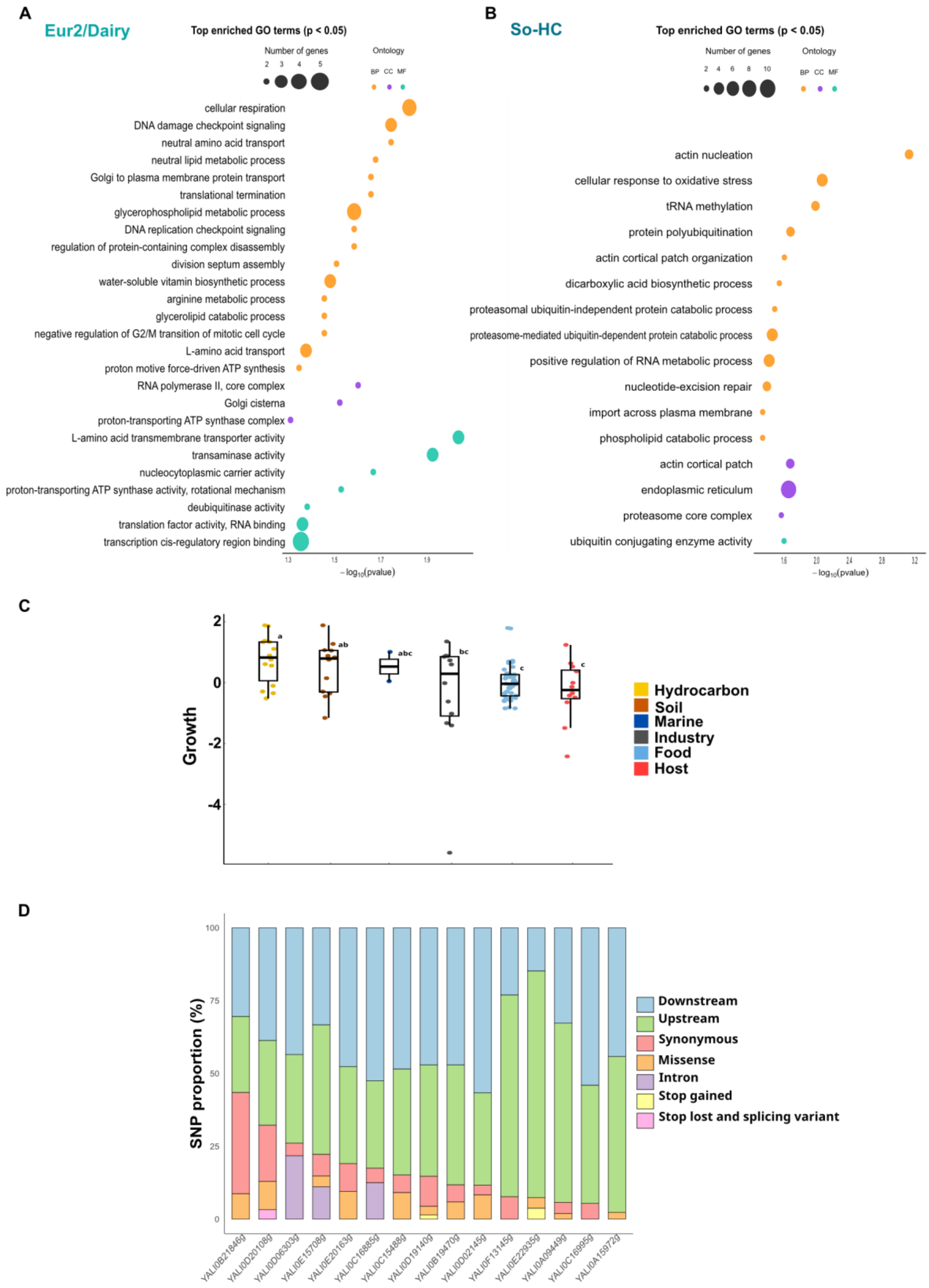

A
Eur2/Dairy
Top enriched GO terms (p < 0.05)
Number of genes
Ontology
BP CC MF
cellular respiration
DNA damage checkpoint signaling
neutral amino acid transport
neutral lipid metabolic process
Golgi to plasma membrane protein transport
translational termination
glycerophospholipid metabolic process
DNA replication checkpoint signaling
regulation of protein-containing complex disassembly
division septum assembly
water-soluble vitamin biosynthetic process
arginine metabolic process
glycerolipid catabolic process
negative regulation of G2/M transition of mitotic cell cycle
L-amino acid transport
proton motive force-driven ATP synthesis
RNA polymerase II, core complex
Golgi cisterna
proton-transporting ATP synthase complex
L-amino acid transmembrane transporter activity
transaminase activity
nucleocytoplasmic carrier activity
proton-transporting ATP synthase activity, rotational mechanism
deubiquitinase activity
translation factor activity, RNA binding
transcription cis-regulatory region binding
−log10(pvalue)
B
So-HC
Top enriched GO terms (p < 0.05)
Number of genes
Ontology
BP CC MF
actin nucleation
cellular response to oxidative stress
tRNA methylation
protein polyubiquitination
actin cortical patch organization
dicarboxylic acid biosynthetic process
proteasomal ubiquitin-independent protein catabolic process
proteasome-mediated ubiquitin-dependent protein catabolic process
positive regulation of RNA metabolic process
nucleotide-excision repair
import across plasma membrane
phospholipid catabolic process
actin cortical patch
endoplasmic reticulum
proteasome core complex
ubiquitin conjugating enzyme activity
−log10(pvalue)
C
Growth
a
ab
abc
bc
c
c
Hydrocarbon
Soil
Marine
Industry
Food
Host
D
SNP proportion (%)
Downstream
Upstream
Synonymous
Missense
Intron
Stop gained
Stop lost and splicing variant
YALI0B21846g
YALI0D20108g
YALI0D06303g
YALI0E15708g
YALI0E20163g
YALI0C16885g
YALI0C15488g
YALI0D19140g
YALI0B19470g
YALI0D02145g
YALI0F13145g
YALI0E22935g
YALI0A09449g
YALI0C16995g
YALI0A15972g

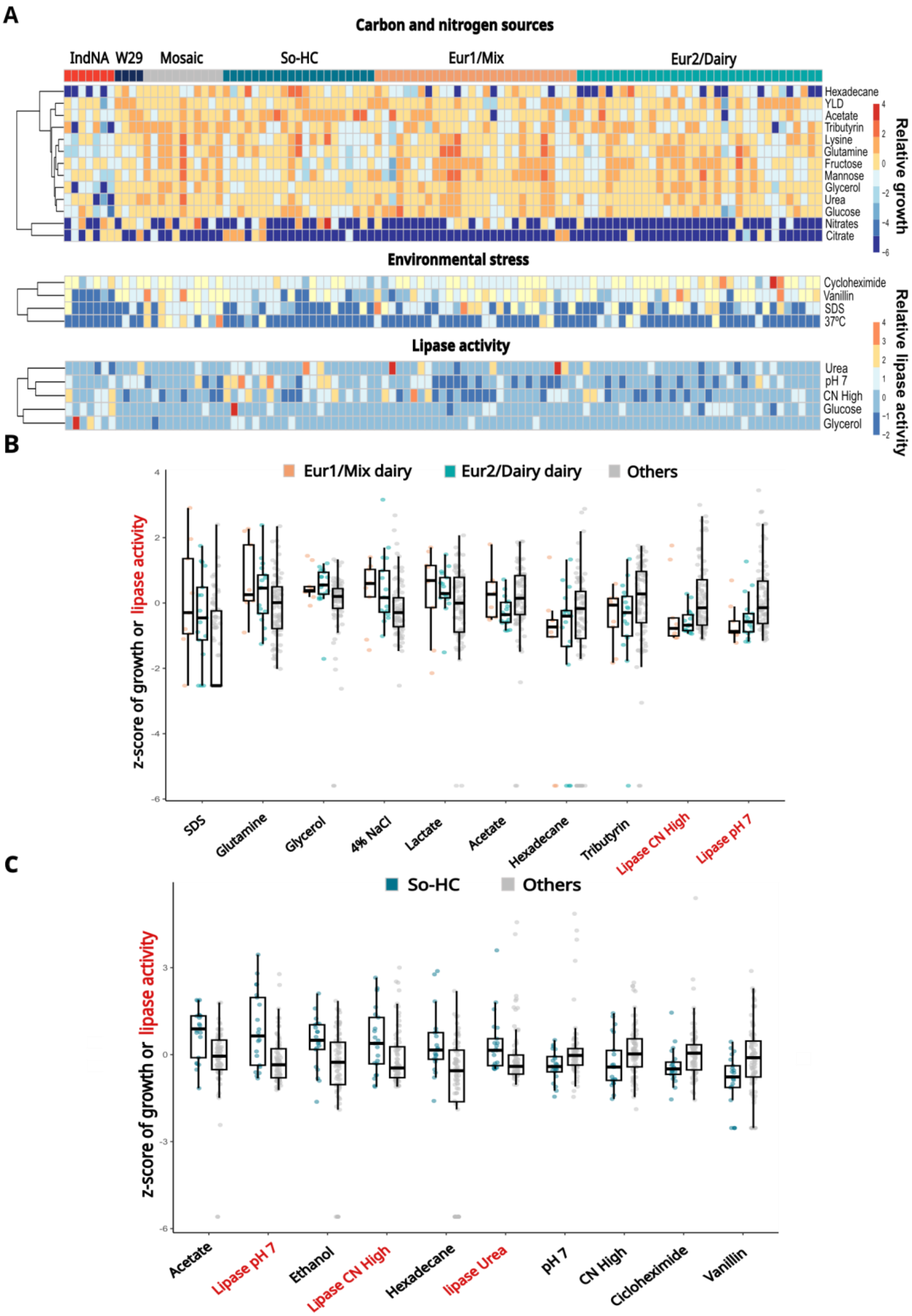
A
Carbon and nitrogen sources
IndNA
W29
Mosaic
So-HC
Eur1/Mix
Eur2/Dairy
Hexadecane
YLD
Acetate
Tributyrin
Lysine
Glutamine
Fructose
Mannose
Glycerol
Urea
Glucose
Nitrates
Citrate
Relative growth
Environmental stress
Cycloheximide
Vanillin
SDS
37°C
Lipase activity
Urea
pH 7
CN High
Glucose
Glycerol
Relative lipase activity
B
Eur1/Mix dairy
Eur2/Dairy dairy
Others
z-score of growth or lipase activity
SDS
Glutamine
Glycerol
4% NaCl
Lactate
Acetate
Hexadecane
Tributyrin
Lipase CN High
Lipase pH 7
C
So-HC
Others
z-score of growth or lipase activity
Acetate
Lipase pH 7
Ethanol
Lipase CN High
Hexadecane
lipase Urea
pH 7
CN High
Cicloheximide
Vanillin
